# A spatial atlas of mitochondrial gene expression reveals dynamic translation hubs and remodeling in stress

**DOI:** 10.1101/2024.08.05.604215

**Authors:** Adam Begeman, John A. Smolka, Ahmad Shami, Tejashree Pradip Waingankar, Samantha C. Lewis

## Abstract

Mitochondrial genome expression is important for cellular bioenergetics. How mitochondrial RNA processing and translation are spatially organized across dynamic mitochondrial networks is not well understood. Here, we report that processed mitochondrial RNAs are consolidated with mitoribosome components into translation hubs distal to either nucleoids or processing granules in human cells. During stress, these hubs are remodeled into translationally repressed mesoscale bodies containing messenger, ribosomal, and double-stranded RNA. We show that the highly conserved helicase SUV3 contributes to the distribution of processed RNA within mitochondrial networks, and that stress bodies form downstream of proteostatic stress in cells lacking SUV3 unwinding activity. We propose that the spatial organization of nascent chain synthesis into discrete domains serves to throttle the flow of genetic information in stress to ensure mitochondrial quality control.

## MAIN TEXT

Mitochondria are endosymbiotic organelles with essential roles in energy production, metabolic regulation, and the innate immune response (*1*). Mitochondria have their own genome encoding 13 respiratory chain complex proteins but rely on nuclear genes to control mitochondrial DNA (mtDNA) replication, transcription, and transcript processing (*2*). Each mammalian cell contains hundreds to thousands of mitochondrial genomes that are individually packaged into complexes termed mitochondrial nucleoids, the units of mtDNA inheritance and sites of transcription (*2*). Constitutive processing granules associate with nucleoids to render the polycistronic mitochondrial RNA into mature messenger, transfer, ribosomal, and non-coding RNAs (*3*). Mitochondrial DNA replication and transcription are thought to be coupled. While most mtDNAs are tightly packaged and likely inaccessible, the sole mitochondrial RNA polymerase, POLRMT, both primes replication and executes processive transcription on a permissible subset of the nucleoid population (*2*). Because each nucleoid contains only 1-2 copies of mtDNA, many nucleoid complexes are distributed throughout the mitochondrial syncytium, where they are asynchronously replicated and transcribed (*4*). In contrast to the nuclear genome, for which DNA replication, transcription, processing and translation occur in distinct compartments, all steps of mitochondrial gene expression co-occur within the innermost compartment of each mitochondrion, the matrix.

The spatial organization of mitochondrial gene expression across dynamic mitochondria that fuse, divide, and are motile in the cytoplasm is not well understood. The core component of the nucleoid complex is mtDNA binding protein TFAM, which has regulatory roles in replication initiation and in transcription via its control of mtDNA compaction, and thus accessibility (*5*, *6*). The RNA processing granules comprise several Fas-activated serine/threonine kinases (FASTK family), as well as G-rich sequence factor 1 protein (GRSF1) which interacts with mitochondrial RNaseP to stimulate primary transcript processing (*3*, *7*, *8*). The leucine-rich pentatricopeptide repeat protein LRPPRC forms a complex with RNA-binding protein SLIRP to bind and stabilize mitochondrial RNAs and is required for their loading to the mitoribosome (*9*). How the processed and polyadenylated transcripts navigate between the processing granules and mitoribosome loading has been obscure, though previous work established that some mitochondria translate more than others within the same cell, suggesting that RNA localization within mitochondrial networks may be regulated to tune electron transport chain (ETC) biogenesis (*10–13*).

Content mixing, facilitated by cycles of membrane fusion and fission, permits the distribution of nascent ETC components, as well as nucleic acids throughout mitochondrial networks. This is particularly important in cells harboring deleterious mtDNA mutations (*14*). Due to its multicopy nature, mutant and wildtype mtDNA often co-exists within cells; complementation of individual organelles via distribution of wildtype gene products from sites of translation confers resilience to ETC dysfunction (*15*). Similarly, distribution of mtDNA gene products can buffer the effects of mtDNA depletion (*16*).

Defects in mtDNA expression cause mitochondrial dysfunction and are linked to cancer, aging and neurodegeneration (*1*). Mitochondrial RNA degradation specifically has emerged as a point of focus (*17*). In homeostasis, the ATP-dependent helicase SUV3 and the mitochondrial PNPase complex function in a linear pathway to unwind and degrade nucleic acids in the matrix, suppressing the persistence of double-stranded RNA that can form by complementarity of H- and L-strand transcripts (*17*, *18*). During stress, accumulation and egress of dsRNA to the cytosol triggers antiviral IFN-1 and pro-inflammatory pathways (*17*, *19*). While both SUV3 and the PNPase complex are required to suppress accumulation of dsRNA in human mitochondria, only PNPase defects lead to mitochondrial leakage and the activation of an inflammatory cascade (*19*, *20*). These findings indicated that while PNPase down-regulation precipitates pro-inflammatory type I IFN responses, it is SUV3 that acts as the upstream gatekeeper of dsRNA accumulation in mitochondria. Consistently, while the PNPase complex is not well conserved between model organisms, SUV3 is highly evolutionarily conserved in sequence and function amongst all eukaryotes, highlighting its essentiality (*21*).

Inborn errors in SUV3 cause neurodegenerative disease in humans despite no evidence of dsRNA release, suggesting an intrinsic mitochondrial stress response upstream of PNPase-dependent organelle permeabilization (*20*). Consistently, a defect in any of multiple steps of mitochondrial gene expression, from mtDNA synthesis, to transcription, to RNA processing, triggers a complex integrated stress response and suppression of mitoribosome translation by unclear mechanism(s) (*22*, *23*). These findings suggest foundational and functional links between the regulation of the mitochondrial central dogma within dynamic networks and quality control of the mitochondrial proteome. Here, we defined how mitochondrial gene expression is spatially organized at sub-organellar scales into regulatory hubs that are amenable to stress-induced remodeling to protect mitochondria during elevated proteostasis burden.

## Results

### Mitochondrial mRNA is excluded from nucleoids and processing granules

Mitochondrial DNA and polycistronic RNA are packaged into distinct nucleoprotein complexes, dedicated to mtDNA synthesis and transcript processing, respectively. However, it is unclear whether the translation of messenger RNAs occurs at defined sites in mitochondrial networks or is coordinated with nucleoids and/or the mitochondrial RNA processing granules (MRGs). Thus, we first sought to directly visualize the spatial distribution of mitochondrial messenger RNAs relative to mtDNA nucleoid complexes or MRGs (**Fig 1A**). We refined a method of fluorescence in situ hybridization (mtRNA-FISH) using fluorophore-conjugated probe sets complementary to processed mRNA, ribosomal RNA, or tRNAs (**Fig S1A**). We validated mitochondrial RNA-FISH signals by confirming their RNaseA-sensitivity and dependence on active transcription by POLRMT (**Fig S1B-C**). We then simultaneously imaged mitochondria, mtDNA, mtRNA and RNA-binding protein GRSF1, a well characterized marker of the total MRG population in mammalian cells, at high spatial resolution, using Airyscan confocal microscopy in IMR90 non-immortalized human fibroblasts. We found that mitochondrial messenger RNAs encoding subunits of Complex I, Complex IV and the mitochondrial ATP Synthase were focally distributed, while in contrast diffuse, ubiquitous mt-tRNA and ribosomal RNA signals marked all mitochondria (**Fig 1B-D; Fig S2**). None of the RNA species we examined colocalized with dsDNA puncta, and linescan analysis indicated that RNA foci were independent of nucleoid positioning along mitochondrial tubules (**Fig 1B-D, far right**). As expected, GRSF1-positive MRGs were evenly distributed among mitochondria and intersected with nucleoids significantly more often than expected by random, as previously reported for other cell types (**Fig 1E-G**). Surprisingly, messenger RNA puncta were nearly twice as abundant as GRSF1 puncta along mitochondria (**Fig 1F**), and the majority did not colocalize with GRSF1 immunofluorescence signals (**Fig 1G**). These observations suggested an order to the distribution of processed mRNAs in mitochondrial networks, beyond their relationship with MRGs.

**Fig. 1.**
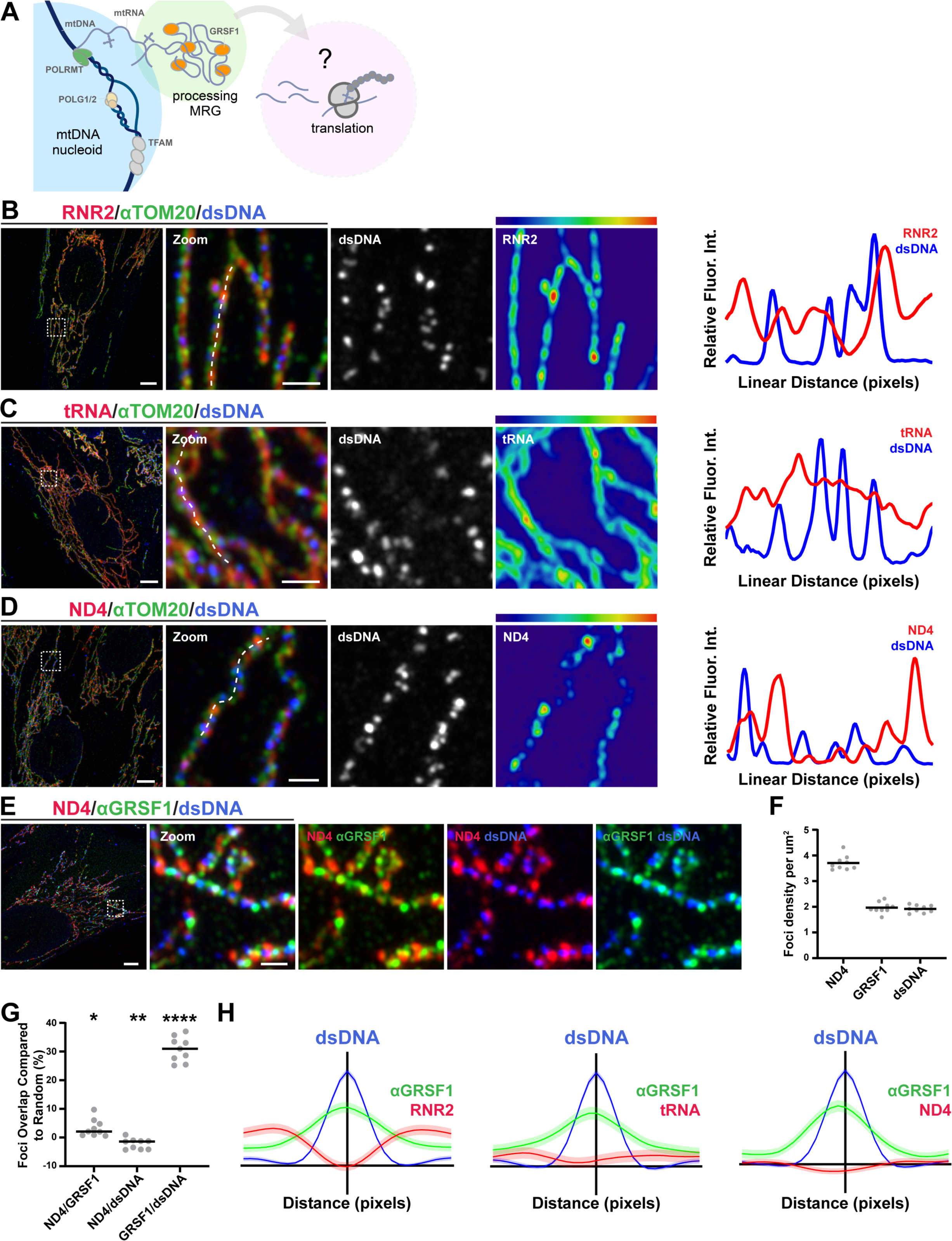
Processed mitochondrial RNA is excluded from nucleoids and MRGs. (A) Overview of mammalian mitochondrial gene expression pathway. (B) Representative images and linescans of fixed IMR90 cells immunolabeled with antibodies against TOM20 (green), dsDNA (blue), and RNA-FISH targeting mitoribosomal component RNR2 (red), (C) mt-transfer RNAs (red), or (D) ND4 messenger RNA (red). Scale bars 5 μm; 1 μm in zoom. (E) IMR90 cell immunolabeled with antibodies against GRSF1 (green), dsDNA (blue), and RNA-FISH targeting ND4 mRNA (red). Scale bars 5 μm; 1 μm in zoom. (F) Comparison of the density of ND4-FISH, anti-GRSF1, and anti-dsDNA foci normalized to mitochondrial area. (G) The frequency of total ND4-FISH, anti-GRSF1, and anti-dsDNA foci that overlap by 20% or more in maximum intensity projections in pairwise comparison. Dotted line represents the frequency of overlap expected by random chance, given the foci density along mitochondrial tubules. (H) Iterative linescan analyses of GRSF1 and RNA localization relative to mtDNA nucleoids marked by anti-dsDNA immunofluorescence and RNA-FISH labeling: RNR2 (n=1392 nucleoids from 21 cells); tRNA (n= 1020 nucleoids from 13 cells); ND4 (n=1211 nucleoids from 26 cells). (****P<0.0001, **P<0.01, *P<0.05, one-sided t-test).

To distinguish mRNA foci from the polycistron we compared RNA-FISH labeling to pulse-labeling of nascent RNA using the click chemistry-compatible nucleoside analog 5-Ethynyluridine (EU) (**Fig S3A-B**) (*24*). GRSF1-positive MRGs colocalized with EU-labeled RNA puncta (**Fig S3C-E**), but not with processed RNA signals reported by FISH.

To rigorously quantify the spatial distributions of nascent and processed RNA across entire mitochondrial networks relative to nucleoids and MRGs, we developed and implemented an image analysis pipeline to systematically map and compare punctate fluorescent signals along filamentous mitochondria at the cellular scale (**Fig S3F-G**). We segmented mitochondria using a machine learning approach, skeletonized them, and extracted linescans to computationally identify peaks of fluorescence intensity in each channel along every mitochondrion. We then used this information from thousands of mitochondria in dozens of cells to generate average fluorescence intensities along a typical linescan segment, in essence, a virtual representation of nucleic acid organization in a typical mitochondrion. We validated our approach by demonstrating that dsDNA and the mtDNA nucleoid marker protein TFAM were highly correlated in these data, as were the EU and GRSF1 intensities (**Fig S3H, left, right)**. We found that dsDNA and GRSF1 intensities were spatially linked to a lesser although still significant degree, consistent with our previous observation (**Fig S3H, middle**). With this tool in hand, we developed a spatial atlas of ribosomal RNA, tRNAs, and mRNA relative to mtDNA and MRGs. Strikingly, RNA-FISH signals not only failed to correlate with nucleoid or MRG markers, but were in fact significantly anti-correlated (**Fig 1H**). These findings indicate that not only are mRNA puncta distinct from nucleoids and MRGs, but they are also surprisingly excluded from those complexes.

We then used CRISPR Cas9 technology to ask whether RNA distribution was dependent on mtDNA copy number by knocking out POLG, the catalytic subunit of the sole mitochondrial DNA polymerase, using three guide RNAs targeting its second exon (**Fig S4A-B**). While the abundance of total mitochondrial RNA scaled to mtDNA nucleoid content (**Fig S4C-D**), ND4-FISH remained punctate, even within the subset of mitochondria that lacked nucleoids altogether (**Fig S4D**). Taken together, we conclude that mitochondria contain focal assemblies of processed RNA that are independent of either the mtDNA nucleoids where mtDNA replication and transcription occur, or MRGs, the nexus of nascent polycistron processing.

### Mitochondrial mRNA marks punctate translation hubs

Mitochondrial translation rates are known to vary among individual mitochondria, though whether there may be a spatial relationship between mitochondrial mRNA distribution and newly synthesized translation products has been unclear (*11*, *13*). Based on our observations of RNA-FISH signals, we hypothesized that mitochondrial mRNA puncta define microscopically visible domains where translation occurs. Thus, we sought to examine the relative localization of mitoribosomes, mRNA, and nascent protein synthesis, taking ND4 as a representative mRNA (**Fig 2A**). We visualized mitoribosome assemblies via indirect immunofluorescence with an antibody against MRPL23, a component of the hydrophobic peptide exit tunnel of the large subunit (mt-LSU) (*25*). Strikingly, MRPL23 signals were punctate, and the majority colocalized with ND4-FISH (**Fig 2B-C; Fig S5A-E**).

**Fig. 2.**
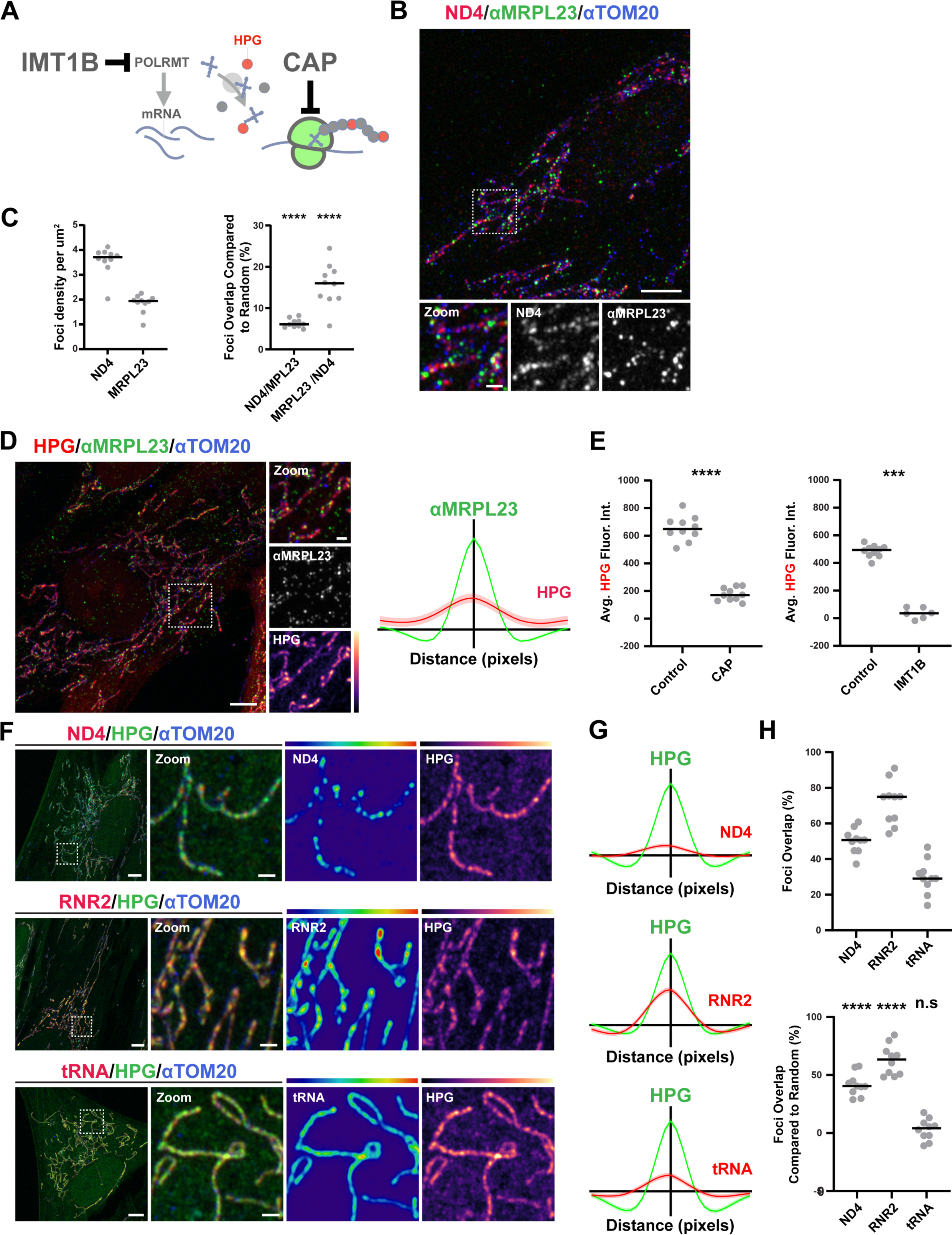
Mitochondrial RNA enrichment marks local translation hubs. (A) Metabolic labeling approach for imaging and manipulating mitochondrial translation. (B) Representative image of fixed IMR90 cells immunolabeled with antibodies against TOM20 (blue), MRPL23 (green), and RNA-FISH targeting ND4 mRNA (red). Scale bars 5 μm; 1 μm in zoom. (C) (Left) Comparison of the density of ND4-FISH and anti-MRPL23 foci normalized to mitochondrial area. (Right) The frequency of ND4-FISH and anti-MRPL23 foci overlap greater than expected by random chance. ****P<0.0001, one-sided t-test. (D) (Left) Representative image of IMR90 cell fixed and immunolabeled to detect MRPL23 (green) and TOM20 (blue) after a 15-minute pulse of 50 μM HPG (red). Scale bars 5 μm; 1 μm in zoom. (Right) Iterative linescan analysis of MRPL23 (n= 4048 from 40 cells) and ND4-FISH fluorescence intensity. (E) (Left) Average HPG fluorescence intensity in segmented mitochondria in control cells versus following 20 min pulse of 50 ug/mL Chloramphenicol (CAP). (Right) Average HPG fluorescence intensity in segmented mitochondria in control cells versus after a 48-hour pulse with 10 uM IMT1B, a mitochondrial RNA polymerase inhibitor. ****P<0.0001, ***P<0.001, Mann-Whitney test. (F) Visualization of HPG fluorescence intensity (green) within mitochondria relative to RNA-FISH (red) targeting ND4 mRNA (top), RNR2 (middle), or mt-tRNAs (bottom). At right, RNA-FISH and HPG are shown color-coded for intensity. Scale bars 5 μm; 1 μm in zoom. (G) Iterative linescans analyses of RNA fluorescence intensities (red) relative to HPG foci (green): ND4 (n=3870 HPG foci from 44 cells); RNR2 (n= 4052 nucleoids from 36 cells); tRNA (n=4522 nucleoids from 41 cells). (H) (top) The proportion of segmented RNA foci that overlap with HPG foci by 20% or more in maximum intensity projections. (Bottom) As above, adjusted relative to the frequency of overlap expected by random chance. ****P<0.0001, one-sided t-test.

We performed translation imaging by pulse-labeling cells with L-HomoPropargylGlycine (HPG), a Methionine analog that is readily recognized by the mitochondrial tRNA^Met^ and incorporated into growing peptide chains and detected using Copper click chemistry (*26*). After a 15-minute pulse, we observed that focal HPG signals colocalized with MRPL23, which was further supported by iterative linescan analysis (**Fig 2D**). HPG labeling was sensitive to the mitoribosome-specific peptidyl-transferase inhibitor chloramphenicol (CAP), as well as the selective POLRMT inhibitor IMT1B (**Fig 2E; Fig S5F**), validating that these observations reflect the output of steady-state gene expression (*27*). Consistent with our earlier observations, HPG-labeled translation domains were not spatially linked with mtDNA nucleoids marked by TFAM, or mtRNA processing granules marked by GRSF1 (**Fig S5G-H**). HPG signals were spatially linked to ND4 puncta, as well as local peaks of RNR2-FISH and tRNA-FISH intensity (**Fig 2F**). In contrast to the relationship between the processed RNAs and nucleoids, ND4-, RNR2- and tRNA-FISH signals correlated well with HPG via iterative linescan (**Fig 2G**). Moreover, the majority of ND4 puncta overlapped with HPG, to an extent significantly above random chance (**Fig 2H**). Taken together, we conclude that mitochondrial protein synthesis occurs in domains or hubs that are marked by mRNA puncta.

### Translation hubs are dynamic and remodeled when mitochondrial fission is defective

We next used a series of pulse-chase experiments to ask whether the translation hubs reported by HPG were dynamic and responsive to network remodeling. We first labeled cells with HPG for 15 minutes; chased in HPG-free medium for 5, 15, 30, or 60 minutes; and then subsequently analyzed the density, size, and average fluorescence intensity of translation hubs after fixation (**Fig 3A**). Consistent with our previous observations, after 5 min of chase, HPG fluorescence intensity was initially restricted to punctate domains (**Fig 3A, top row**). Quantification of HPG signals during the series of extended chase experiments revealed that HPG signals progressively and dramatically decreased with lengthening chase time (**Fig 3B-D**), consistent with nascent chain degradation and/or the distribution away from those sites throughout mitochondria. This observation was important, because it indicated that in fibroblasts, translation hubs serve as point sources for ETC component proteins.

**Fig. 3.**
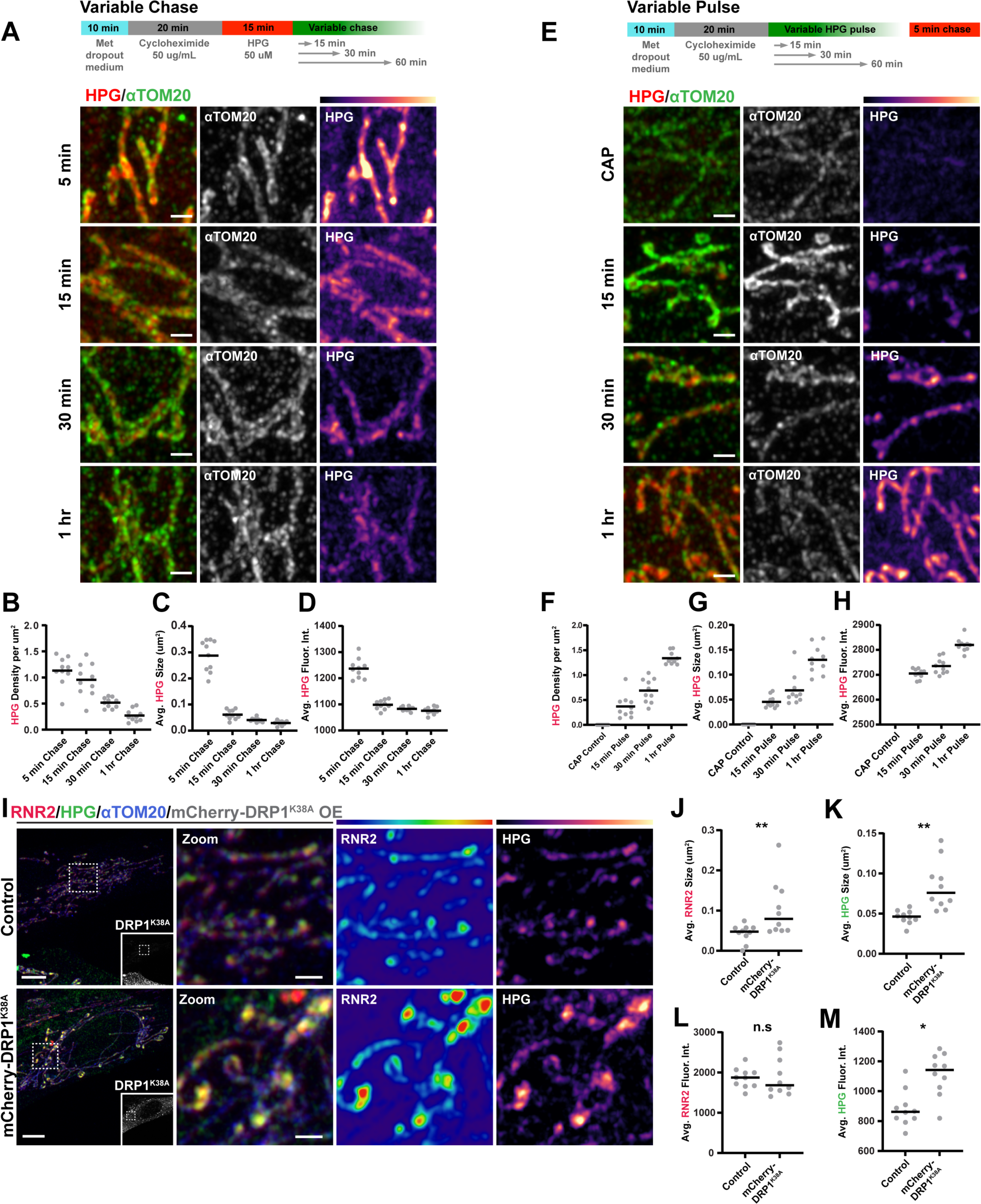
Mitochondrial translation hubs are dynamic. (A) (Top) Overview of HPG labeling time course with variable *chase* times. (Bottom) Representative images of IMR90 cells pulse labeled for 15 minutes with HPG and chased with unlabeled methionine for 5, 15, 30, or 60 minutes. Scale bar 1 μm. (B) Density of thresholded HPG domains normalized to mitochondrial area in each condition. (C) Average size of HPG-labeled domains in each condition. (D) Average above-threshold fluorescence intensity of HPG-labeled domains in each condition. (E) (Top) Overview of HPG labeling time course with variable *pulse* times. (Bottom) Representative images of IMR90 cells pulse labeled for 15, 30, or 60 minutes with HPG followed by a constant chase in unlabeled methionine for 5 minutes, relative to control cells incubated in HPG for 15 minutes concurrent with 50 uM CAP. Scale bar 1 μm. (F-H) Quantification of the number of thresholded HPG objects per segmented mitochondrial area in each condition. (I) Representative images of IMR90 cells that were transiently transfected with mCherry-DRP1^K38A^ (greyscale), pulse labeled for 15 minutes with 50 uM HPG (green), immunolabeled with an antibody against TOM20 (blue), and RNR2-FISH (red). Scale bar 5 μm; 1μm in zoom. (J) Average size of thresholded RNR2 signal intensity. (K) Average size of thresholded HPG signal intensity. (L) Average fluorescence intensity of RNR2 signals per thresholded mitochondrion. (M) Average fluorescence intensity of HPG per thresholded mitochondrion. (**P<0.01,*P<0.05, Mann-Whitney Test).

We then sought to assess whether all mitochondria are capable of translation under our experimental conditions. We pulsed cells with HPG for 15, 30, or 60 minutes while holding the chase constant at 5 minutes and again measured features of HPG labeling among mitochondria (**Fig 3E**). We found that the number of HPG-labeled domains per mitochondrion increased in a manner directly proportional to pulse length (**Fig 3F**), though heterogeneity across the entire mitochondrial network persisted (**Fig S5I**). The size and fluorescence intensity of individual HPG hotspots increased as well (**Fig 3G-H**). These observations indicate that local translation shapes protein distribution within mitochondria, as peptide synthesis occurs at discrete sites and protein products then diffuse along the inner membrane, are distributed to other mitochondrial fragments via membrane dynamics, or are degraded.

Mitochondrial content mixing is in large part dependent on fission, fusion, and motility dynamics governed by dynamin family guanosine triphosphatases (GTPases) (*28*). DRP1, a cytosolic <u>d</u>ynamin-related <u>p</u>rotein, forms helical scission assemblies around mitochondria mediated by its interactions with receptors in the outer membrane and by the close apposition of membrane contact sites with other organelles (*29*). To test the idea that compartmentalized gene expression in translation hubs shapes the flow of genetic information in mitochondrial networks at the cellular level, we next examined the role of mitochondrial fusion-fission cycles. We used transient overexpression of mCherry-tagged DRP1^K38A^, a dominant mutation in the GTP-binding pocket that disrupts GTP hydrolysis, to decrease mitochondrial fission rate and assess the impact on the distribution of nascent translation products (**Fig 3I**) (*29*, *30*). Transient overexpression of mCherry-DRP1^K38A^ caused significant mitochondrial elongation as compared with control cells as predicted (**Fig S5J**), above and beyond baseline mitochondrial elongation caused by cycloheximide pre-treatment. We found that suppressing mitochondrial fission led to increased size and fluorescence intensity of both RNA- and HPG-enriched translation hubs in a subset of mitochondria, consistent with defective content mixing (**Fig 3J-M**). Thus, when mitochondrial fission is perturbed, newly synthesized ETC proteins labeled by HPG fail to be distributed, reducing network homogenization. These data confirm that dynamic translation hubs and fusion/fission cycles shape the distribution of nascent peptides across mitochondrial networks.

### Mitochondrial RNA is remodeled into translationally repressed liquid-like mesoscale bodies during stress

We hypothesized that the organization of the matrix into dynamic translation hubs could serve to facilitate a rapid and local translational response to perturbation. Previous work has shown that pathogenic variants that disrupt mitochondrial protein synthesis converge on activation of a complex integrated stress response (ISR), characterized in part by shutoff of mitoribosome translation and accumulation of aberrant double-stranded mitochondrial RNA (*22*, *23*, *31*, *32*). During ISR activation, release of mitochondrial dsRNA into the cytosol triggers a pro-inflammatory transcriptional response, because the nucleic acid is recognized as foreign (*17*). Similarly, outer membrane permeabilization elicited by mitochondrial poisons also permits nucleic acid egress, which contributes to an mtDNA-triggered innate immune response (*33–35*). Given that context, we sought a means by which we could test whether translation hubs may be remodeled to facilitate translational inhibition during dsRNA accumulation - without triggering inflammatory cascades that might confound our imaging-based approach.

SUV3 is an essential mitochondrial ATP-dependent RNA helicase that unwinds both double-stranded RNA (dsRNA) and RNA:DNA hybrids; it is also important for the nucleolytic degradation of aberrant, proteotoxic dsRNAs by the mitochondrial PNPase complex (**Fig 4A**) (*17*, *36*). In human cells, SUV3 silencing causes dsRNA accumulation (*36*). Importantly, multiple studies in cells and in patients with inborn errors in SUV3 have shown that it functions upstream of the PnPase complex in a linear pathway, such that loss of SUV3 function alone is insufficient for dsRNA release to the cytosol or detectable activation of innate immunity (*19*, *20*). Thus, we used depletion and transient overexpression of wildtype and mutant SUV3 isoforms to examine a potential role for translation hubs in mediating mitochondrial stress responses to dsRNA accumulation.

**Fig. 4.**
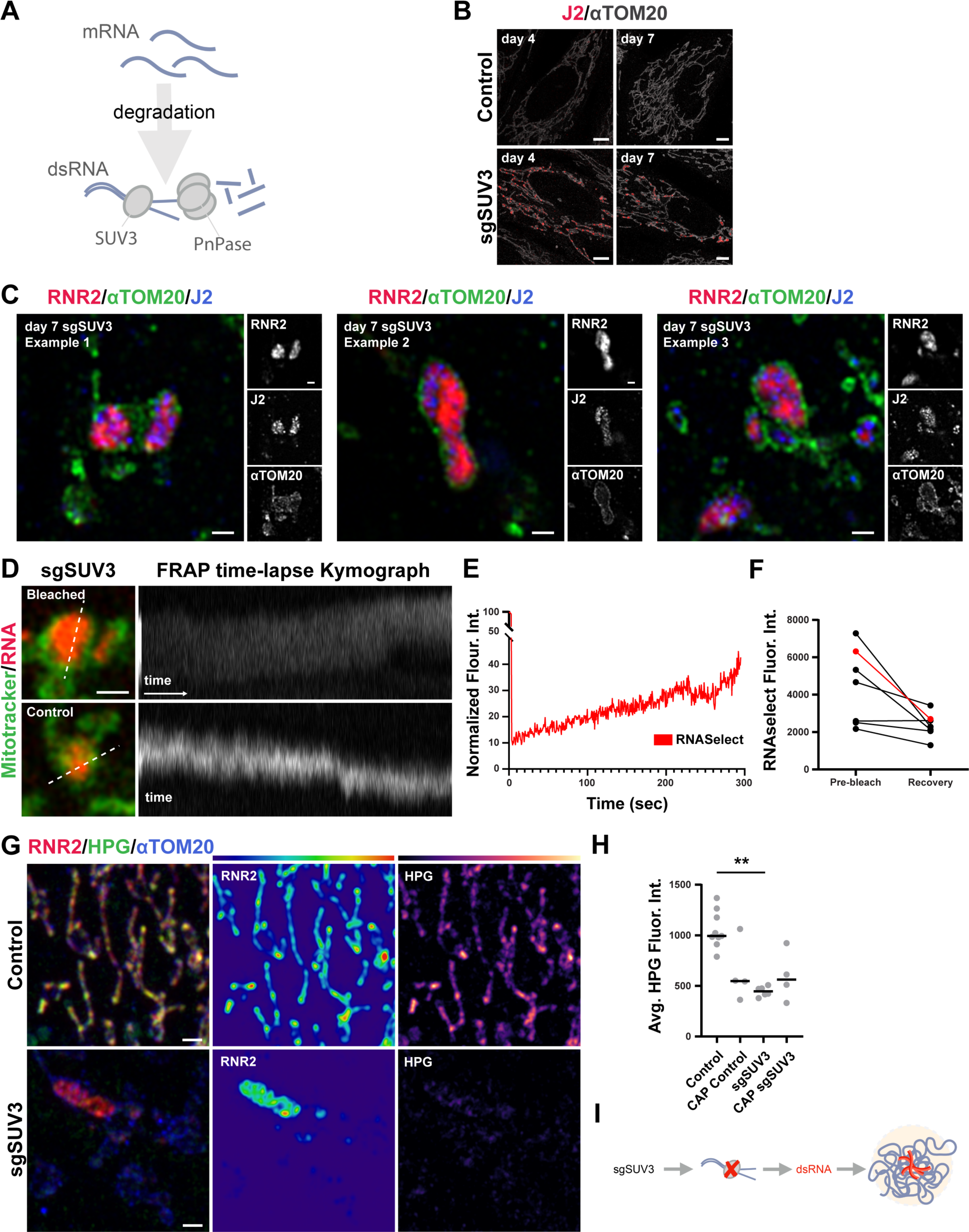
Mitochondrial RNAs remodel into translationally repressed mesoscale bodies during stress. (A) Schematic of mitochondrial transcript unwinding and degradation by SUV3 and PnPase. (B) Representative images of cells immunolabeled for TOM20 (greyscale) and dsRNA (J2; red) at 4 days versus 7 days of SUV3 depletion. Scale bar 5 μm. (C) Three representative images of cells fixed and immunolabeled with antibodies against dsRNA (J2; blue), TOM20 (green), and RNA-FISH targeting RNR2 (red). Scale bar 1 μm; 1 μm in zoom. (D) Representative images and kymographs from live cells labeled with Mitotracker Deep Red (green) and SYTO RNASelect (red), during photobleach and recovery (Top) or unbleached control (Bottom). Scale bar 1 μm. (E) FRAP intensity for representative bleach and recovery of SYTO RNASelect over 300 seconds. (F) Comparison of pre-bleach and recovery fluorescence intensities for 7 mesoscale bodies from 7 cells over a 300 second interval. (G) Representative image of control (Top) and SUV3-depleted (Bottom) cells pulse-labeled with 50 uM HPG for 15 min (green), fixed and immunolabeled to detect TOM20 (blue), and RNA-FISH against RNR2 (red). Scale bar 1 μm. (H) Average HPG fluorescence intensity in segmented mitochondria in the presence or absence of 50 ug/mL CAP. (**P<0.01, Kruskal-Wallis test (**P<0.01), followed by Dunn’s multiple comparisons. (I) Diagram of proposed impact of SUV3 depletion on dsRNA accumulation and single-stranded RNA reorganization.

Transient overexpression of SUV3^WT^-HA in control IMR90 cells revealed its localization to a multitude of discrete puncta coincident with a subset of endogenous GRSF1 foci at MRGs (**Fig S6A**), consistent with a role in suppressing hybridization between complementary endogenous RNAs. We next depleted SUV3 from cells by CRISPR Cas9 technology using three guide RNAs targeting exon 1 and examined mitochondrial dsRNA levels over a 7-day time course using indirect immunofluorescence with the anti-dsRNA antibody J2. While dsRNA was undetectable in control cells, we found that cells depleted of SUV3 continuously accumulated dsRNA, which coalesced into distinct foci within a subset of malformed, swollen mitochondria (**Fig 4B**). We then examined ssRNA distribution, finding that RNR2-FISH signals were no longer distributed throughout mitochondrial networks, but collapsed into distended boli while rendering large areas of the mitochondrial network devoid of ribosomal RNA (**Fig S6B, top**). To verify that this phenotype was specifically due to the lack of SUV3 helicase activity, we reintroduced either SUV3^WT^-HA or SUV3^G207V^-HA, a dominant, catalytically dead mutant allele, into the cells (**Fig S6B-C**). Transient overexpression of SUV3^WT^-HA, but not SUV3^G207V^-HA, rescued RNR2 localization, indicating that loss of SUV3 function not only leads to dsRNA accumulation, but remodeling of ssRNA distribution as well.

We next asked whether the accumulated dsRNA may seed the formation of the larger ssRNA-enriched boli. We examined fixed cells depleted of SUV3 by indirect immunofluorescence to simultaneously visualize TOM20, dsRNA via J2, and RNR2-FISH (**Fig 4C**). Indeed, we observed that RNR2-FISH signals had coalesced into boli surrounding the dsRNA explaining the distended appearance of the mitochondrial membranes. Co-labeling of cells with ND4-FISH and immunodetection of dsDNA revealed the re-organization of mRNA into the enlarged boli as well (**Fig S7**). Despite this, nucleoids remained distributed suggesting that mitochondrial DNA and RNA may be positioned in mitochondria by distinct mechanisms.

Previous studies have posited that mitochondrial ribonucleo-protein complexes may exhibit properties of phase-separated condensates (*37–39*). Given our observations that processed RNAs are excluded from nucleoids, and the striking remodeling of RNA in cells depleted of SUV3, we next sought to examine the dynamics of RNA in live cells. We labeled RNA in live IMR90 cells with the vital dye SYTO RNASelect and co-stained with the vital dye Mitotracker. Consistent with our observations in fixed cells, RNASelect labeled many discrete puncta in control cells, while it labeled prevalent mesoscale structures when SUV3 was limiting (**Fig 4D; Fig S8A**), that maintained their membrane potential as reported by both Mitotracker and tetramethylrhodamine ethyl ester (TMRE) staining (**Fig S8B**).

We next asked whether these mesoscale bodies exhibited properties of liquid-like membraneless RNA bodies, similar to Balbiani bodies or RNP granules induced by viral infection, by assessing RNA dynamics and propensity for content exchange by time-lapse microscopy and fluorescence recovery after photobleaching (FRAP) (*37*, *40*). RNA boli labeled by RNASelect-labeled exhibited dynamic, fluctuating morphologies over time, though they persisted as discrete domains within mitochondria, with infrequent fusion or fission (**Fig 4D; Fig S8A, bottom**). Indeed, we also observed that multiple RNASelect-labeled domains would often co-persist within the same mitochondrion. We employed FRAP to determine whether these structures exchanged contents, finding that RNASelect intensity recovered on average to 40% of the pre-bleach intensity after background correction over a time period of 5 minutes (**Fig 4E-F**). This was less and slower recovery than previously reported for MRGs, but much more recovery than seen for solid mitochondrial aggregates composed exclusively of protein previously described in yeasts (*37*, *41*). Additional biochemical studies are needed to determine whether these structures could be bona fide biocondensates. Given their dynamic nature, stress-specific context, and evidence of content exchange, we will refer to them here as “mitochondrial RNA stress bodies” (MSB).

To determine the relationship between MSB formation and mitochondrial translation, we next used RNA-FISH and HPG pulse-labeling to assess protein synthesis in cells with the MSB phenotype. Relative to control cells, SUV3 depletion led to a near total loss of detectable HPG incorporation, concurrent with MSB formation (**Fig 4G**). Indeed, the intensity of HPG labeling in SUV3-depleted cells was comparable to control cells incubated with chloramphenicol (CAP) (**Fig 4H**). These findings demonstrate critical functional dependencies between dsRNA accumulation, the spatial organization of RNA in the matrix, and steady state mitoribosome translation, by which MSB formation is linked to translational inhibition (**Fig 4I**).

### Mesoscale body formation is linked to proteostasis

The reorganization of RNA into MSBs prompted us to investigate the fates of the pre-existing MRGs during that process. We visualized the localization of endogenous GRSF1 in IMR90 cells labeled with RNR2-FISH by indirect immunofluorescence during SUV3 depletion (**Fig S9**). Unlike control cells, we observed a significant increase in GRSF1 colocalization with processed RNA in MSBs (**Fig S9**). In addition, we noted that while MSBs were marked by GRSF1, separate small GRSF1 puncta remained distributed throughout mitochondrial networks that did not colocalize with RNR2-FISH signals, consistent with a continued role in binding to and processing polycistronic RNA at MRGs and similar to the distribution we noted for nucleoids. Thus, while MSB’s contain dsRNA, ssRNA, and a typical MRG protein, these structures are spatially distinct from MRGs and remain so over time.

Finally, we considered whether MSB formation is a response to proteotoxicity downstream of defective mtRNA processing or e.g. dsRNA accumulation. We reasoned that, if MSBs form in response to an RNA processing defect alone, suppression of mitoribosome translation would have little effect on their formation or dynamics, as the RNA would still be produced. In contrast, if quality control were regulated at the protein level, only cells that actually translate aberrant messages would trigger MSB formation. Thus, we asked whether the preemptive arrest of mitoribosomes by CAP could suppress MSB formation. In distinct experiments, we performed a CAP pulse-chase analysis in cells either before (**Fig 5A**) or after (**Fig 5B**) transfection with sgSUV3-Cas9 RNP complexes, and subsequently analyzed the size and fluorescence intensity of MSBs. Preemptive mitoribosome arrest via CAP dramatically reduced the size of FISH-labeled MSBs relative to sgSUV3 cells treated with DMSO only (**Fig 5B**). In contrast, turning off the mitoribosome after SUV3 depletion failed to suppress MSB size or fluorescence intensity. To further define the contribution of proteotoxicity to MSB formation, we asked whether dsRNA accumulation and MSB formation are separable. We incubated cells with CAP, induced SUV3 depletion, then fixed them and detected dsRNA and RNR2-FISH. Consistent with our previous observation, CAP treatment suppressed the MSB phenotype; moreover, the cells still accumulated dsRNA (**Fig 5C**). This experiment demonstrated that MSB formation is a response to a proteostatic stress, as it relies on active translation by mitoribosomes. Consistently, we found that, in the absence of CAP, in early stages of MSB formation the dsRNA is apparent before RNR2 remodeling into boli (**Fig 5D**). Taken together, these data suggest a model in which homeostatic mitochondrial translation hubs enriched in processed RNA are remodeled into MSB structures in coordination with translational inhibition to protect proteostasis (**Fig 5E**).

**Fig. 5.**
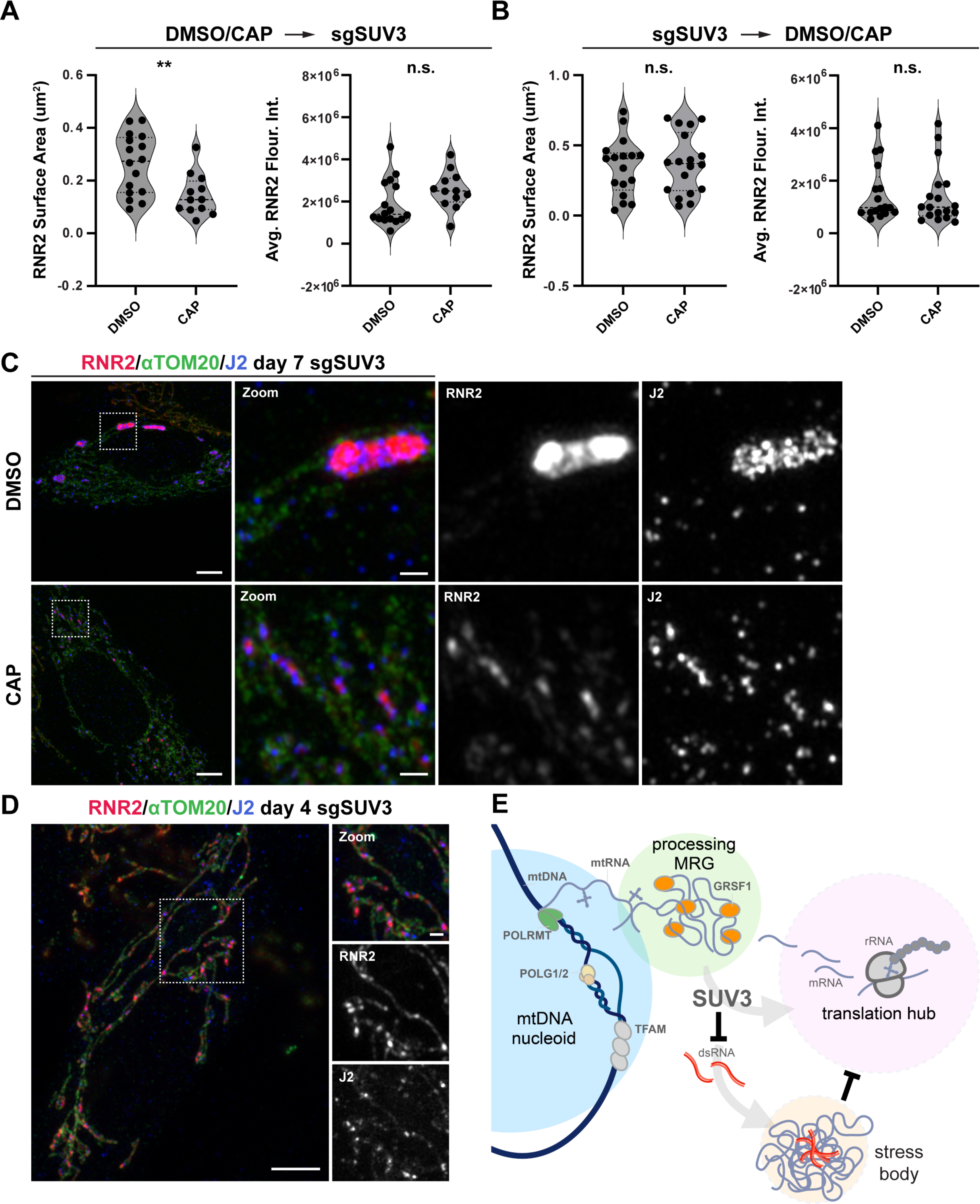
Mitochondrial RNA remodeling in stress is proteoprotective. (A) Average size and fluorescence intensity of segmented RNR2-FISH signals in IMR90 cells when SUV3 depletion is induced in cells already translationally repressed by incubation in 50 ug/mL CAP. (B) Average size and fluorescence intensity of segmented RNR2-FISH signals in IMR90 cells incubated in 50 ug/mL CAP 7 days after transfection with sgSUV3 RNP complexes. (**P<0.01, Mann-Whitney Test). (C) Representative images of control (Top) and 50 uM CAP-treated (Bottom) cells 7 days after transfection with sgSUV3 RNP complexes,, fixed and immunolabeled to detect dsRNA (J2; blue), TOM20 (green), and RNA-FISH against RNR2 (red). Scale bar 5 μm; 1μm in zoom. (D) Representative image of cells 4 days after transfection with sgSUV3 RNP complexes, fixed and immunolabeled to detect dsRNA (J2; blue), TOM20 (green), and RNR2-FISH against RNR2 (red). Scale bar 5 μm; 1μm in zoom. (E) Proposed model for organization of mitochondrial gene expression into translational hubs. SUV3-dependent dsRNA accumulation, translational suppression, and mesoscale body formation protect the mitochondrial proteome during stress.

## Discussion

Our data indicate that within human mitochondrial networks, processed RNAs are translated in ribonucleo-protein hubs distinct from mtDNA nucleoids or MRGs, and these hubs are remodeled in response to RNA pre-processing stress, concurrent with suppression of mitoribosome translation. We propose that the spatial organization of mtDNA nucleoids, MRGs, and processed messages at these sites provides a means of throttling the flow of genetic information to ensure quality control of the mitochondrial proteome. Such a process of remodeling and translational suppression may be particularly important for mitochondria, as once expressed, core components of the electron transport chain complexes are long-lived (*42*).

It will be important to determine the fundamental molecular mechanisms by which mitoribosome shutdown occurs. Our analyses highlight the importance of the sub-mitochondrial organization of gene expression in homeostasis, and how that organization is remodeled in stress via sequestration of mtRNA and proteins within context-specific mitochondrial RNA stress bodies. Liquid-like condensates have important functions in regulating gene expression in the cell nucleus and cytoplasm, where they provide a means to locally concentrate sets of proteins and RNAs for regulation in 4-dimensions (*40*, *43–45*). Both mitochondrial nucleoids and MRGs have been suggested to have biophysical properties of condensates; how those properties may contribute to mtDNA or RNA integrity remains to be discerned (*37*, *38*).

Compartmentalization of biochemical processes is a unifying principle in biology. In eukaryotes, the nuclear envelope provides a means to control the entry and egress of transcription machinery and products, which may be processed and are ultimately translated in distinct and spatially segregated compartments. In the cytosol, ribonucleoprotein granules, such as P-bodies that form during animal development, and yeast stress granules, serve as hubs to organize post-transcriptional regulation of gene expression. These well-characterized bodies consolidate translationally repressed mRNAs to regulate where and when gene expression occurs, which is particularly important in stress. Within mitochondria, the mitochondrial genome, immature polycistronic transcripts, and processed RNAs in various stages share the milieu of the matrix compartment. In this capacity, remodeling of the matrix into the structures described here could function to modulate the accessibility of mitoribosome loading or accessory factors to mRNAs that would normally shuttle between these compartments, tuning production of ETC proteins to sub-cellular cues. This could be particularly important in highly polarized cells such as neurons, in which mitochondrial functions may need to be specialized for the cell soma, dendrite, and axonal compartments (*46*). Given that SUV3 loss-of-function causes neurodegenerative disease in humans, it will be important to investigate whether MSBs exhibit further features of biocondensates and to refine our understanding of the conditions under which they form, as constitutive MSBs may constitute a form of pathological inclusion (*20*, *47*, *48*). Beyond SUV3, LRPPRC and/or SLIRP are likely to mediate both translation hub activity and MSB formation, given their key roles in protecting mitochondrial RNAs from degradation as well as in mitoribosome loading.

Our findings connect the spatial organization of the steps of mitochondrial genome expression in the matrix to the kinetics of mitoribosome translation and overall mitochondrial network morphometrics. This connection has implications for understanding the cellular pathology and complex stress responses underlying metabolic dysregulation. Developing approaches that can suppress or modulate the flow of genetic information in mitochondria may hold promise in the treatment of rare human diseases caused by defects in mitochondrial gene expression, as well as in the context of ISR activation during cancer and neurodegeneration.

## Materials and Methods

### Plasmids

The mCherry-DRP1^K38A^ plasmid was generated via QuickChange mutagenesis from mCherry-Drp1, a gift from Gia Voeltz (Addgene #49152) (*30*). To generate SUV3-HA and SUV3G207V-HA, the human SUV3 cDNA was synthesized (Twist Bioscience) with or without the G207V amino acid substitution, including an HA tag appended to the C-terminus of the sequence, and cloned into the pTwist CMV-driven mammalian expression vector.

### Mammalian cell growth, transfection, and vital dyes

Human IMR90 cells (ATCC #CCL-186) were grown in high-glucose Dulbecco’s Modified Eagle’s medium (DMEM) supplemented with 10% heat inactivated fetal bovine serum (FBS) and 1% penicillin/streptomycin at 37 degrees Celsius in a humidified 5% CO^2^ chamber. Prior to imaging, cells were seeded onto glass-bottom 35 mm dishes (Mattek) and cultured for 24-48 hours. Transient plasmid transfections were performed using Lipofectamine 2000 according to the manufacturer protocol (Thermo Fisher), and imaged 24 hours later unless otherwise noted. For live cell imaging, 1 mL of conditioned cell media was removed and saved and then, 50 nM Mitotracker Deep Red (Thermo Fisher), 500 nM Tetramethylrhodamine, Ethyl Ester (TMRE) (Thermo Fisher), or 5 uM SYTO RNAselect (Thermo Fisher) was added directly to the dish for 30 minutes and then replaced with the reserved conditioned media just prior to imaging.

### Cell fixation, antibodies, and immunofluorescence

Cells were seeded as described above. Cells were then fixed in pre-warmed (37 degrees Celsius) 4% paraformaldehyde solution (PFA) in DPBS for 30 minutes at room temperature while protected from light. Glass-bottom dishes were then gently washed with room temperature DPBS and cells were permeabilized in 0.1% TritionX-100 diluted in DPBS for 20 minutes. Dishes were washed with TBST blocking buffer (TBS pH 7.5, 0.1% Tween-20) containing 1% bovine serum albumin (BSA), and primary antibodies were added at 1:1000 dilution in the same buffer and incubated for 1 hour at room temperature, or overnight at 4 degrees Celsius. Dishes were then rinsed with blocking buffer and a solution containing secondary antibodies at 1:2000 dilution was added for 1 hour at room temperature. Dishes were rinsed with the blocking buffer and imaged in DPBS at room temperature. We employed the following antibodies to detect endogenous proteins and nucleic acids: rabbit anti-TOM20 (Proteintech, 11802-1-AP), mouse anti-TOM20 (SantaCruz Biotechnology, SC17764), mouse anti-dsDNA (Abcam, ab27156), rabbit anti-GRSF1 (Sigma, HPA036984), rabbit anti-TFAM (Abcam, ab176558), rabbit anti-MRPL23 (Sigma, HPA050406), mouse anti-DRP1 (Abnova, H00010059), rabbit anti-HA (Invitrogen, 71-5500), mouse anti-J2 (Sigma, MABE1134), donkey anti-rabbit AlexaFluor Plus 488 highly cross-adsorbed conjugate (ThermoFisher, A32790), goat anti-mouse AlexaFluor 405 conjugate (ThermoFisher, A31553), donkey anti-mouse AlexaFluor Plus 405 highly cross-adsorbed conjugate (ThermoFisher, A48257), goat anti-rabbit AlexaFluor 405 conjugate (ThermoFisher, A31556), goat anti-mouse AlexaFluor Plus 647 highly cross-adsorbed conjugate (ThermoFisher, A32728).

### Mitochondrial RNA fluorescence *in situ* hybridization (FISH)

Labeling of mitochondrial RNA via fluorescence *in situ* hybridization was performed using a modified version of the Stellaris RNA-FISH protocol (Biosearch Technologies). All probe sequences are available in **Supplementary Table 1**. Cells were seeded onto plates as described above. When FISH was combined with copper click chemistry to detect nascent proteins, then HPG labeling and the click reaction were performed prior to the FISH protocol and the cells were fixed and immunolabeled as described above. Otherwise, the cells were fixed in prewarmed (37 degrees Celsius) 3% PFA, 1.5% glutaraldehyde in DPBS for 10 minutes at room temperature. The fixation reaction was then quenched with a solution containing a 1:10 dilution of 1 M glycine in DPBS and washed with DPBS. To quench auto-fluorescence associated with glutaraldehyde, dishes were then incubated with 10 mg/mL sodium borohydride diluted in DPBS for 5 minutes at room temperature. Cells were then washed with DPBS and permeabilized in 0.1% TritionX-100, 1 uM dithiothreitol (DTT), 0.1% SDS in DPBS for 20 minutes at room temperature. Following permeabilization, dishes were washed with DPBS and incubated in Stellaris wash buffer A, prepared according to manufacturer’s protocol (Biosearch Technologies), for 5 minutes. Wash buffer A was then aspirated and 200 uL of RNA-FISH probe set containing Stellaris hybridization buffer, prepared according to the manufacturer’s protocol, was then added directly to the center of the glass bottom dish. Dishes were then incubated in a hybridization chamber at 37 degrees Celsius for 4 hours. Following incubation, the hybridization buffer was removed by aspiration and dishes were incubated in Stellaris wash buffer A at 37 degrees Celsius for 30 minutes. Dishes were then washed with Stellaris wash buffer B, immunolabeled, and imaged as described above. For IMT1B control 10 uM final concentration was used for 96 hours prior to labeling and for RNaseA 100 ug/mL was used at room temperature for 1 hour after fixation and permeabilization but prior to FISH labeling.

### EU Click reaction for labeling of nascent RNA

Labeling of nascent mitochondrial RNA was performed using ethynyl-uridine (EU) incorporation and resulting click reaction. Cells were first seeded and cultured as described above. On the day of the experiment, 1 mL of conditioned media was removed and set aside, then 1 uL of 1 mM triptolide was then added directly to cells in the imaging dish at a final concentration of 1 uM and samples were incubated for 30 minutes at 37 degrees Celsius. Next 1 uL of 500 mM EU was added to the sample media to a final concentration of 500 uM, and incubated for 1 hour. EU- and triptolide-containing media was then replaced with the reserved1 mL of conditioned media and incubated for a final 5 minutes. Following fixation and permeabilization as described above, EU incorporation was detected via Click-iT Plus Edu Alexa Fluor 647 imaging kit (REF) according to manufacturer’s instructions, and then immunolabeled and imaged as described above.

### HPG Click reaction for labeling of nascent peptide synthesis

Metabolic labeling of proteinactive mitochondrial translation was performed using a modified version of the manufacturer’s Click-IT Homopropargylglycine (HPG) protocol (Thermo Fisher). Cells were first seeded and cultured as described above. Cells were then washed with warm (37 C) PBS and then incubated in methionine-free DMEM for 10 minutes before the addition of drugs to suppress either cytosolic protein translation (50 ug/mL Cycloheximide) or mitochondrial protein translation (50 ug/mL Chloramphenicol). Cells were then pulsed with HPG (50 uM final concentration) added directly to the drug containing methionine-free DMEM and incubated for 5-60 minutes as indicated in the text. Cells were then chased with methionine free DMEM lacking HPG as described. Following fixation and permeabilization as described above, samples then underwent a click chemistry reaction to render HPG fluorescent via the covalent addition of AlexaFluor probes following the manufacturer’s protocol (Thermo Fisher).

### Live Cell Imaging and FRAP

For FRAP assays, cells were seeded on poly-d-lysine-coated glass bottom 35 mm imaging dishes and cluttered for 1 to 2 days in normal growth conditions and then labeled with vital dyes as described above. The 148 (0.268um^2) pixel bleaching ROIs were placed on representative RNAselect bolus in sgSUV3 condition and 5 time points were taken prior to bleaching to establish baseline. A different cell and frame of view was chosen for each FRAP experiment and samples exchanged after 2 hours of imaging. Recovery was monitored for 500 frames at 0.59 seconds per frame for a total of ∼300 seconds.

### Microscopy and image acquisition

All images were acquired using a Zeiss LSM 980 with Airyscan 2 laser scanning confocal microscope, equipped with 405, 588, 561, and 639 nanometer laser lines and Fast Airyscan detector array. Images were acquired using an inverted 63x/1.4 NA oil objective. All live imaging was done in a humidified chamber at 37 C and in the presence of 5% CO^2^. Airyscan processing and maximum intensity image projection was performed using Zeiss ZEN Blue software version 3.7 (Carl Zeiss). Image brightness and/or contrast were linearly adjusted in ZEN Blue or FIJI (*49*).

### Image Analysis

FIJI, Arivis, and Python were used. All Z stack images were 3D Airyscan processed and maximum intensity projections were generated using Zeiss ZEN Blue software version 3.7 (Carl Zeiss) and saved as czi files. These projections were then converted to Arivis sis file format using the Arivis SIS batch converter (ver 4.1.0) for subsequent analysis in Arivis Vision4D (ver 4.1). All graphs and other visualization of quantification was created using Graphpad Prism 10 (ver 10.1.0) All Arivis segmentation pipelines, machine learning trainings and models, as well as all custom analysis code for FIJI, and Python available at https://github.com/TheLewisLab.

#### Segmentation

Segmentation of the mitochondrial network was achieved using the Arivis machine learning image trainer to design a machine learning model using the Fluorescence and EM Robust training dataset for all channels in which mitochondria were labeled. The trainer was trained using, on average, 3 representative images from the image set, classifying mitochondria and background signal until sufficient segmentation of the mitochondrial network was achieved and artifacts minimized. For Mitotracker deep red, RNR2 and tRNA signals, an intensity threshold segmenter was used to define mitochondrial objects. The resulting segments were then filtered for size, using a 0.100 - 0.200 um^2^ cutoff.

For punctate signals (ND4, GRSF1, dsDNA, TFAM, MRPL23), the Arivis blob finder method was used. For more continuous signal, domains of enrichment were defined using the intensity-based threshold segmenter. All objects were then filtered by their proximity to the mitochondrial network segment using the Arivis compartmentalization function with a 60% object intersection cutoff. Object intersections within the mitochondria were also determined using the Arivis compartmentalization function with a 20% cutoff. Relevant object information, such as area, intensity measurements, or intersection data, was then exported as a csv for downstream analysis. The resulting mitochondrial network segmentation was used as an image mask and exported from Arivis as OME TIFFs for downstream analysis.

For HPG variable pulse and variable chase intensity thresholding, different intensity values were chosen for the variable pulse (2500 gray value) and variable chase (1000 gray value) due to the differing dynamic range in captures between datasets.

#### Object intersection analysis

To determine the mitochondrial object intersection percentages above expected by random chance, object intersection percentages of segmented objects were compared to simulated data using the average sizes, number, and mitochondrial area data extracted from Arivis segmentation above. In short, a Monte Carlo simulation was performed 100 times using the extracted object data to determine how often the two objects would overlap at least 20% by area given their respective sizes, abundances, and the compartment area of the mitochondrial network. This simulation was run for each frame of view and compared to the extracted object intersection values generated from the Arivis segmentation pipeline.

#### Automated linescan extraction

To quantify fluorescence intensity along linescans from skeletonized mitochondrial networks, we generated a custom Fiji macro, as follows. In short, mitochondria in micrographs were masked as described above and the mask was skeletonized using a binary image threshold via the Skeletonize 3D Fiji Package (*50*). The ‘Analyze Skeleton’ feature was then applied to systematically map object branch points, endpoints, and junctions within the mitochondrial network. This function was performed iteratively 5 times, each time pruning branches of less than 8 pixels in length to remove artificial branches generated during the skeletonization. After the skeleton was pruned of artificial branches the junction pixels were removed as well and the Analyze skeleton getShortestPathPoints() function used to generate polylines across the mitochondrial network. These lines were then added to the ROI manager, and using the Fiji ProfilePlot function, fluorescence intensity linescans in all channels were generated for each line. The resulting csv files were then used for downstream analysis.

#### Linescan analysis and peak calling

Peak calling and analysis of the fluorescence intensity linescans was performed using custom Python code available at https://github.com/TheLewisLab. Peaks of fluorescence intensity were called using the scipy find_peaks function for each extracted linescan (*51*). For each peak called in a given fluorescence channel, the 0.5 um on either side was extracted and averaged to form the average signal over a given fluorescence peak. Random average line scans were generated by randomly selecting a peak position across the same line scan dataset and averaging the 1 um surrounding that point. Averages, confidence intervals, and relevant statistics were performed using the native python packages.

### Generation and validation of CRISPR KOs

Guide RNA sequences used to target POLG and SUV3 are available in **Supplementary Table 2**. To generate knockout cells via CRISPR Cas9 gene editing, we implemented a modified version of the Lipofectamine CRISPRMax Cas9 Reagent lipofectamine protocol (ThermoFisher). Cells were transfected with Cas9 ribonucleoparticles pre-complexed with synthetic guide RNAs (Synthego) generated bySynthego Gene Knockout Kit made up of 2NLS-Cas9 and validated synthetic multi guide RNAs. Briefly, cells were seeded to 15-20% density the day prior to transfection in 6 well plates. To prepare the transfection mixture, 2 uL of 20 uM NLS-Cas9 were complexed with 4 uL of 10 uM sgRNAs in 14 uL Optimem (Invitrogen) for a final volume of 20 uL, then mixed. In parallel, 10 uL of Cas9 Plus reagent was diluted into 70 uL of Optimem, mixed, and subsequently added to the complexed Cas9/sgRNA complexes. For transfection, 6 uL of the Lipofectamine CRISPRMax Cas9 Reagent was diluted into 94 uL of Optimem, mixed, and added to the Cas9/sgRNA/Cas9 Plus solution. The mixture was allowed to incubate at room temperature for 10-minute incubation at room temperature before added dropwise to the plated cells. The media was replaced 24 hours later.

## Acknowledgements

The authors thank the members of the Lewis laboratory, as well as Drs. James Hurley, Gary Karpen, and Michael Rape for helpful comments.

## Author Contributions

Conceptualization: AB, SL

Methodology: AB, JS, SL

Investigation: AB, AS, TW, JS, SL

Visualization: AB, SL

Supervision: SL

Writing—original draft: AB, SL

Writing—review & editing: AB, SL

## Data and Materials Availability

All data are available in the main text or the supplementary materials. All code is available on the Lewis laboratory Github site, as described in Materials and Methods. All data are available on request.

## References

1. A. Suomalainen, J. Nunnari, Mitochondria at the crossroads of health and disease. Cell 187, 2601–2627 (2024).

2. C. M. Gustafsson, M. Falkenberg, N.-G. Larsson, Maintenance and Expression of Mammalian Mitochondrial DNA. Annu. Rev. Biochem. 85, 133–160 (2016).

3. H. Antonicka, F. Sasarman, T. Nishimura, V. Paupe, E. A. Shoubridge, The Mitochondrial RNA-Binding Protein GRSF1 Localizes to RNA Granules and Is Required for Posttranscriptional Mitochondrial Gene Expression. Cell Metab. 17, 386–398 (2013).

4. S. C. Lewis, L. F. Uchiyama, J. Nunnari, ER-mitochondria contacts couple mtDNA synthesis with mitochondrial division in human cells. Science 353, aaf5549 (2016).

5. N.-G. Larsson, J. Wang, H. Wilhelmsson, A. Oldfors, P. Rustin, M. Lewandoski, G. S. Barsh, D. A. Clayton, Mitochondrial transcription factor A is necessary for mtDNA maintance and embryogenesis in mice. Nat. Genet. 18, 231–236 (1998).

6. N. A. Bonekamp, M. Jiang, E. Motori, R. G. Villegas, C. Koolmeister, I. Atanassov, A. Mesaros, C. B. Park, N.-G. Larsson, High levels of TFAM repress mammalian mitochondrial DNA transcription in vivo. Life Sci. Alliance 4 (2021).

7. A. A. Jourdain, M. Koppen, M. Wydro, C. D. Rodley, R. N. Lightowlers, Z. M. Chrzanowska-Lightowlers, J.-C. Martinou, GRSF1 Regulates RNA Processing in Mitochondrial RNA Granules. Cell Metab. 17, 399–410 (2013).

8. A. Ohkubo, L. V. Haute, D. L. Rudler, M. Stentenbach, F. A. Steiner, O. Rackham, M. Minczuk, A. Filipovska, J.-C. Martinou, The FASTK family proteins fine-tune mitochondrial RNA processing. PLOS Genet. 17, e1009873 (2021).

9. B. Ruzzenente, M. D. Metodiev, A. Wredenberg, A. Bratic, C. B. Park, Y. Cámara, D. Milenkovic, V. Zickermann, R. Wibom, K. Hultenby, H. Erdjument-Bromage, P. Tempst, U. Brandt, J. B. Stewart, C. M. Gustafsson, N.-G. Larsson, LRPPRC is necessary for polyadenylation and coordination of translation of mitochondrial mRNAs. EMBO J. 31, 443–456 (2012).

10. L. Chatre, M. Ricchetti, Large heterogeneity of mitochondrial DNA transcription and initiation of replication exposed by single-cell imaging. J. Cell Sci. 126, 914–926 (2013).

11. M. Zorkau, C. A. Albus, R. Berlinguer-Palmini, Z. M. A. Chrzanowska-Lightowlers, R. N. Lightowlers, High-resolution imaging reveals compartmentalization of mitochondrial protein synthesis in cultured human cells. Proc. Natl. Acad. Sci. 118, e2008778118 (2021).

12. C. Estell, E. Stamatidou, S. El-Messeiry, A. Hamilton, In situ imaging of mitochondrial translation shows weak correlation with nucleoid DNA intensity and no suppression during mitosis. J. Cell Sci. 130, 4193–4199 (2017).

13. R. Yousefi, E. F. Fornasiero, L. Cyganek, J. Montoya, S. Jakobs, S. O. Rizzoli, P. Rehling, D. Pacheu-Grau, Monitoring mitochondrial translation in living cells. EMBO Rep. 22, e51635 (2021).

14. S. Vidoni, C. Zanna, M. Rugolo, E. Sarzi, G. Lenaers, Why Mitochondria Must Fuse to Maintain Their Genome Integrity. Antioxid. Redox Signal. 19, 379–388 (2013).

15. K. Nakada, K. Inoue, T. Ono, K. Isobe, A. Ogura, Y.-I. Goto, I. Nonaka, J.-I. Hayashi, Inter-mitochondrial complementation: Mitochondria-specific system preventing mice from expression of disease phenotypes by mutant mtDNA. Nat. Med. 7, 934–940 (2001).

16. G. Elachouri, S. Vidoni, C. Zanna, A. Pattyn, H. Boukhaddaoui, K. Gaget, P. Yu-Wai-Man, G. Gasparre, E. Sarzi, C. Delettre, A. Olichon, D. Loiseau, P. Reynier, P. F. Chinnery, A. Rotig, V. Carelli, C. P. Hamel, M. Rugolo, G. Lenaers, OPA1 links human mitochondrial genome maintenance to mtDNA replication and distribution. Genome Res. 21, 12–20 (2011).

17. A. Dhir, S. Dhir, L. S. Borowski, L. Jimenez, M. Teitell, A. Rötig, Y. J. Crow, G. I. Rice, D. Duffy, C. Tamby, T. Nojima, A. Munnich, M. Schiff, C. R. de Almeida, J. Rehwinkel, A. Dziembowski, R. J. Szczesny, N. J. Proudfoot, Mitochondrial double-stranded RNA triggers antiviral signalling in humans. Nature 560, 238–242 (2018).

18. R. J. Szczesny, L. S. Borowski, L. K. Brzezniak, A. Dmochowska, K. Gewartowski, E. Bartnik, P. P. Stepien, Human mitochondrial RNA turnover caught in flagranti: involvement of hSuv3p helicase in RNA surveillance. Nucleic Acids Res. 38, 279– 298 (2010).

19. M. R. Krieger, M. Abrahamian, K. L. He, S. Atamdede, H. Hakimjavadi, M. Momcilovic, D. Ostrow, S. D. Maggo, Y. P. Tsang, X. Gai, G. F. Chanfreau, D. B. Shackelford, M. A. Teitell, C. M. Koehler, Trafficking of mitochondrial double-stranded RNA from mitochondria to the cytosol. Life Sci. Alliance 7 (2024).

20. S. L. van Esveld, R. J. Rodenburg, F. Al-Murshedi, E. Al-Ajmi, S. Al-Zuhaibi, M. A. Huynen, J. N. Spelbrink, Mitochondrial RNA processing defect caused by a mutation in two siblings with a novel neurodegenerative syndrome. J. Inherit. Metab. Dis. 45, 292–307 (2022).

21. R. J. Szczesny, L. S. Borowski, M. Malecki, M. A. Wojcik, P. P. Stepien, P. Golik, RNA Degradation in Yeast and Human Mitochondria. Biochim. Biophys. Acta BBA - Gene Regul. Mech. 1819, 1027–1034 (2012).

22. M. M. Foged, E. Recazens, S. Chollet, M. Lisci, G. E. Allen, B. Zinshteyn, D. Boutguetait, C. Münch, V. K. Mootha, A. A. Jourdain, Cytosolic N6AMT1-dependent translation supports mitochondrial RNA processing. bioRxiv [Preprint] (2024). 10.1101/2024.07.02.601698.

23. C. B. Jackson, A. Marmyleva, R. Awadhpersad, G. Monteuuis, T. Mito, N. Zamboni, T. Tatsuta, A. E. Vincent, L. Wang, T. Langer, C. J. Carroll, A. Suomalainen, De novo serine biosynthesis is protective in mitochondrial disease. bioRxiv [Preprint] (2023). 10.1101/2023.03.23.533952.

24. C. Y. Jao, A. Salic, Exploring RNA transcription and turnover in vivo by using click chemistry. Proc. Natl. Acad. Sci. 105, 15779–15784 (2008).

25. E. Lavdovskaia, E. Hanitsch, A. Linden, M. Pašen, V. Challa, Y. Horokhovskyi, H. P. Roetschke, F. Nadler, L. Welp, E. Steube, M. Heinrichs, M. M.-Q. Mai, H. Urlaub, J. Liepe, R. Richter-Dennerlein, A roadmap for ribosome assembly in human mitochondria. Nat. Struct. Mol. Biol., 1–11 (2024).

26. Y. Kimura, H. Saito, T. Osaki, Y. Ikegami, T. Wakigawa, Y. Ikeuchi, S. Iwasaki, Mito-FUNCAT-FACS reveals cellular heterogeneity in mitochondrial translation. RNA 28, 895–904 (2022).

27. N. A. Bonekamp, B. Peter, H. S. Hillen, A. Felser, T. Bergbrede, A. Choidas, M. Horn, A. Unger, R. Di Lucrezia, I. Atanassov, X. Li, U. Koch, S. Menninger, J. Boros, P. Habenberger, P. Giavalisco, P. Cramer, M. S. Denzel, P. Nussbaumer, B. Klebl, M. Falkenberg, C. M. Gustafsson, N.-G. Larsson, Small-molecule inhibitors of human mitochondrial DNA transcription. Nature 588, 712–716 (2020).

28. B. Westermann, Mitochondrial fusion and fission in cell life and death. Nat. Rev. Mol. Cell Biol. 11, 872–884 (2010).

29. E. Smirnova, L. Griparic, D.-L. Shurland, A. M. van der Bliek, Dynamin-related Protein Drp1 Is Required for Mitochondrial Division in Mammalian Cells. Mol. Biol. Cell 12, 2245–2256 (2001).

30. J. R. Friedman, L. L. Lackner, M. West, J. R. DiBenedetto, J. Nunnari, G. K. Voeltz, ER Tubules Mark Sites of Mitochondrial Division. Science 334, 358–362 (2011).

31. I. Hochberg, L. A. M. Demain, J. Richer, K. Thompson, J. E. Urquhart, A. Rea, W. Pagarkar, A. Rodríguez-Palmero, A. Schlüter, E. Verdura, A. Pujol, P. Quijada-Fraile, A. Amberger, A. J. Deutschmann, S. Demetz, M. Gillespie, I. A. Belyantseva, H. J. McMillan, M. Barzik, G. M. Beaman, R. Motha, K. Y. Ng, J. O’Sullivan, S. G. Williams, S. S. Bhaskar, I. R. Lawrence, E. M. Jenkinson, J. L. Zambonin, Z. Blumenfeld, S. Yalonetsky, S. Oerum, W. Rossmanith, W. W. Yue, J. Zschocke, K. J. Munro, B. J. Battersby, T. B. Friedman, R. W. Taylor, R. T. O’Keefe, W. G. Newman, Bi-allelic variants in the mitochondrial RNase P subunit PRORP cause mitochondrial tRNA processing defects and pleiotropic multisystem presentations. Am. J. Hum. Genet. 108, 2195–2204 (2021).

32. G. Wang, E. Shimada, C. M. Koehler, M. A. Teitell, PNPASE and RNA trafficking into mitochondria. Biochim. Biophys. Acta BBA - Gene Regul. Mech. 1819, 998– 1007 (2012).

33. K. McArthur, L. W. Whitehead, J. M. Heddleston, L. Li, B. S. Padman, V. Oorschot, N. D. Geoghegan, S. Chappaz, S. Davidson, H. San Chin, R. M. Lane, M. Dramicanin, T. L. Saunders, C. Sugiana, R. Lessene, L. D. Osellame, T.-L. Chew, G. Dewson, M. Lazarou, G. Ramm, G. Lessene, M. T. Ryan, K. L. Rogers, M. F. van Delft, B. T. Kile, BAK/BAX macropores facilitate mitochondrial herniation and mtDNA efflux during apoptosis. Science 359, eaao6047 (2018).

34. M. Tigano, D. C. Vargas, S. Tremblay-Belzile, Y. Fu, A. Sfeir, Nuclear sensing of breaks in mitochondrial DNA enhances immune surveillance. Nature 591, 477–481 (2021).

35. A. P. West, G. S. Shadel, Mitochondrial DNA in innate immune responses and inflammatory pathology. Nat. Rev. Immunol. 17, 363–375 (2017).

36. L. S. Borowski, A. Dziembowski, M. S. Hejnowicz, P. P. Stepien, R. J. Szczesny, Human mitochondrial RNA decay mediated by PNPase–hSuv3 complex takes place in distinct foci. Nucleic Acids Res. 41, 1223–1240 (2013).

37. T. Rey, S. Zaganelli, E. Cuillery, E. Vartholomaiou, M. Croisier, J.-C. Martinou, S. Manley, Mitochondrial RNA granules are fluid condensates positioned by membrane dynamics. Nat. Cell Biol. 22, 1180–1186 (2020).

38. M. Feric, A. Sarfallah, F. Dar, D. Temiakov, R. V. Pappu, T. Misteli, Mesoscale structure–function relationships in mitochondrial transcriptional condensates. Proc. Natl. Acad. Sci. 119, e2207303119 (2022).

39. Q. Long, Y. Zhou, H. Wu, S. Du, M. Hu, J. Qi, W. Li, J. Guo, Y. Wu, L. Yang, G. Xiang, L. Wang, S. Ye, J. Wen, H. Mao, J. Wang, H. Zhao, W.-Y. Chan, J. Liu, Y. Chen, P. Li, X. Liu, Phase separation drives the self-assembly of mitochondrial nucleoids for transcriptional modulation. Nat. Struct. Mol. Biol. 28, 900–908 (2021).

40. C. Roden, A. S. Gladfelter, RNA contributions to the form and function of biomolecular condensates. Nat. Rev. Mol. Cell Biol. 22, 183–195 (2021).

41. L. Ruan, J. T. McNamara, X. Zhang, A. C.-C. Chang, J. Zhu, Y. Dong, G. Sun, A. Peterson, C. H. Na, R. Li, Solid-phase inclusion as a mechanism for regulating unfolded proteins in the mitochondrial matrix. Sci. Adv. 6, eabc7288 (2020).

42. S. Krishna, R. Arrojo e Drigo, J. S. Capitanio, R. Ramachandra, M. Ellisman, M. W. Hetzer, Identification of long-lived proteins in the mitochondria reveals increased stability of the electron transport chain. Dev. Cell 56, 2952–2965.e9 (2021).

43. P. A. Sharp, A. K. Chakraborty, J. E. Henninger, R. A. Young, RNA in formation and regulation of transcriptional condensates. RNA 28, 52–57 (2022).

44. S. F. Banani, H. O. Lee, A. A. Hyman, M. K. Rosen, Biomolecular condensates: organizers of cellular biochemistry. Nat. Rev. Mol. Cell Biol. 18, 285–298 (2017).

45. Y. Shin, C. P. Brangwynne, Liquid phase condensation in cell physiology and disease. Science 357, eaaf4382 (2017).

46. G. Pekkurnaz, X. Wang, Mitochondrial heterogeneity and homeostasis through the lens of a neuron. Nat. Metab. 4, 802–812 (2022).

47. A. Zbinden, M. Pérez-Berlanga, P. De Rossi, M. Polymenidou, Phase Separation and Neurodegenerative Diseases: A Disturbance in the Force. Dev. Cell 55, 45–68 (2020).

48. S. Spannl, M. Tereshchenko, G. J. Mastromarco, S. J. Ihn, H. O. Lee, Biomolecular condensates in neurodegeneration and cancer. Traffic 20, 890–911 (2019).

49. J. Schindelin, I. Arganda-Carreras, E. Frise, V. Kaynig, M. Longair, T. Pietzsch, S. Preibisch, C. Rueden, S. Saalfeld, B. Schmid, J.-Y. Tinevez, D. J. White, V. Hartenstein, K. Eliceiri, P. Tomancak, A. Cardona, Fiji: an open-source platform for biological-image analysis. Nat. Methods 9, 676–682 (2012).

50. I. Arganda-Carreras, R. Fernández-González, A. Muñoz-Barrutia, C. Ortiz-De-Solorzano, 3D reconstruction of histological sections: Application to mammary gland tissue. Microsc. Res. Tech. 73, 1019–1029 (2010).

51. P. Virtanen, R. Gommers, T. E. Oliphant, M. Haberland, T. Reddy, D. Cournapeau, E. Burovski, P. Peterson, W. Weckesser, J. Bright, S. J. van der Walt, M. Brett, J. Wilson, K. J. Millman, N. Mayorov, A. R. J. Nelson, E. Jones, R. Kern, E. Larson, C. J. Carey, İ. Polat, Y. Feng, E. W. Moore, J. VanderPlas, D. Laxalde, J. Perktold, R. Cimrman, I. Henriksen, E. A. Quintero, C. R. Harris, A. M. Archibald, A. H. Ribeiro, F. Pedregosa, P. van Mulbregt, SciPy 1.0: fundamental algorithms for scientific computing in Python. Nat. Methods 17, 261–272 (2020).

