## Supplemental Figures S1-9 for "A spatial atlas of mitochondrial gene expression reveals dynamic translation hubs and remodeling in stress"

Adam Begeman *et al.*

**This PDF file includes:**

Figs. S1 to S9

**Other Supplementary Materials for this manuscript include the following:**

Tables S1 to S2

Figure S1

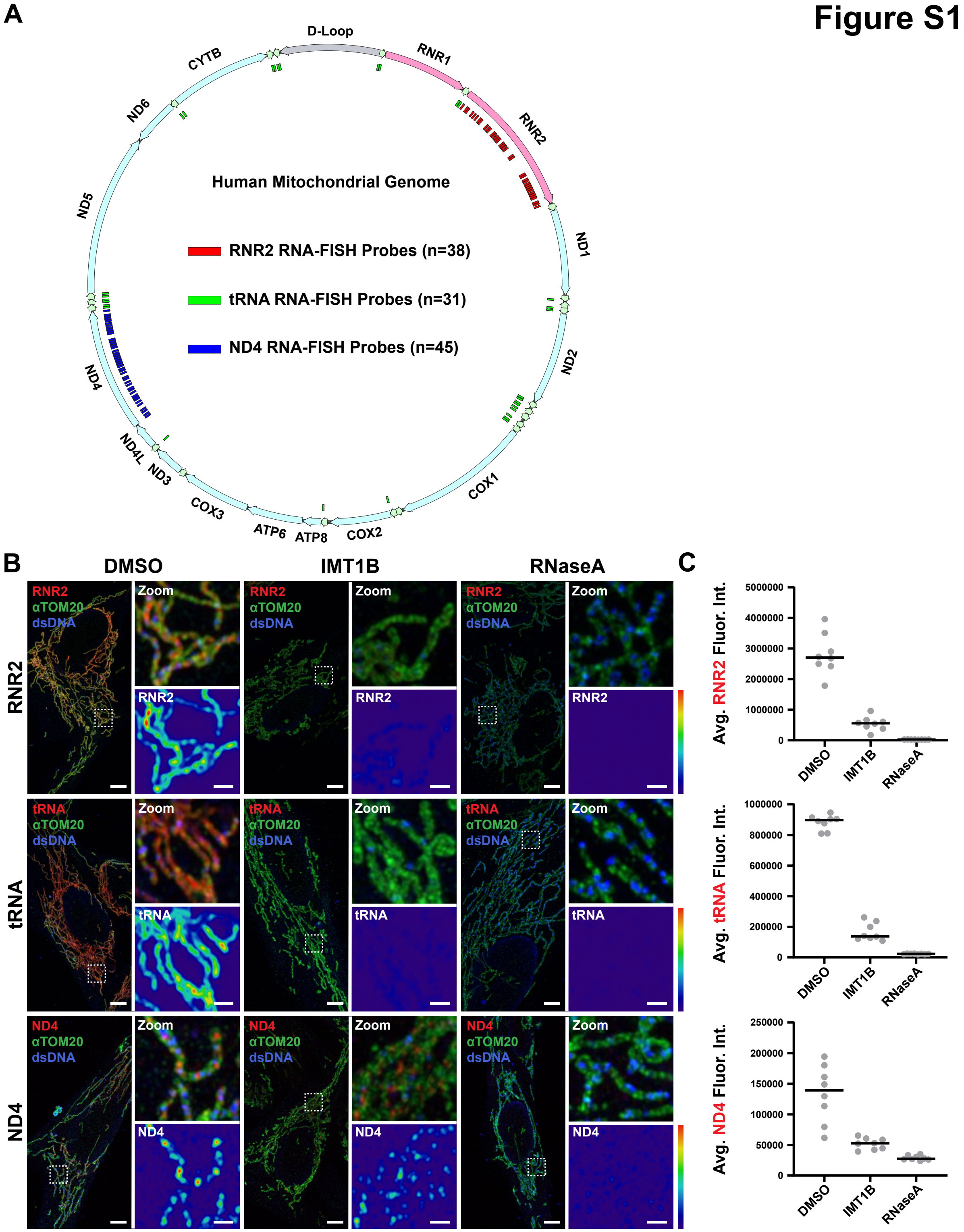
A
D-Loop
RNR1
RNR2
ND1
ND2
COX1
COX2
ATP8
ATP6
COX3
ND3
ND4L
ND4
ND5
ND6
CYTB
Human Mitochondrial Genome
RNR2 RNA-FISH Probes (n=38)
tRNA RNA-FISH Probes (n=31)
ND4 RNA-FISH Probes (n=45)
B
DMSO
IMT1B
RNaseA
RNR2
tRNA
ND4
αTOM20
dsDNA
Zoom
C
Avg. RNR2 Fluor. Int.
Avg. tRNA Fluor. Int.
Avg. ND4 Fluor. Int.
5000000
4000000
3000000
2000000
1000000
0
1000000
800000
600000
400000
200000
0
250000
200000
150000
100000
50000
0
DMSO
IMT1B
RNaseA

**Fig. S1. Specifically visualizing mitochondrial DNA encoded RNAs via RNA-FISH.** (A) Schematic of where the RNA-FISH probes are located on each mitochondrial RNA. (B) Representative images of fixed IMR90 cells treated with DMSO (left), 10 uM IMT1B for 96 hours (middle), or 100 ug/mL RNaseA for 1 hour prior to FISH labeling(right) immunolabeled for TOM20 (green) and dsDNA (blue), and mtRNA-FISH against either RNR2 (top), tRNAs (middle), or ND4 (bottom) (red). Scale bars 5 μm; 1 μm in zoom. (C) Quantification of average RNA intensity per mitochondrial area in each condition for each RNA in (B).

### Figure S2

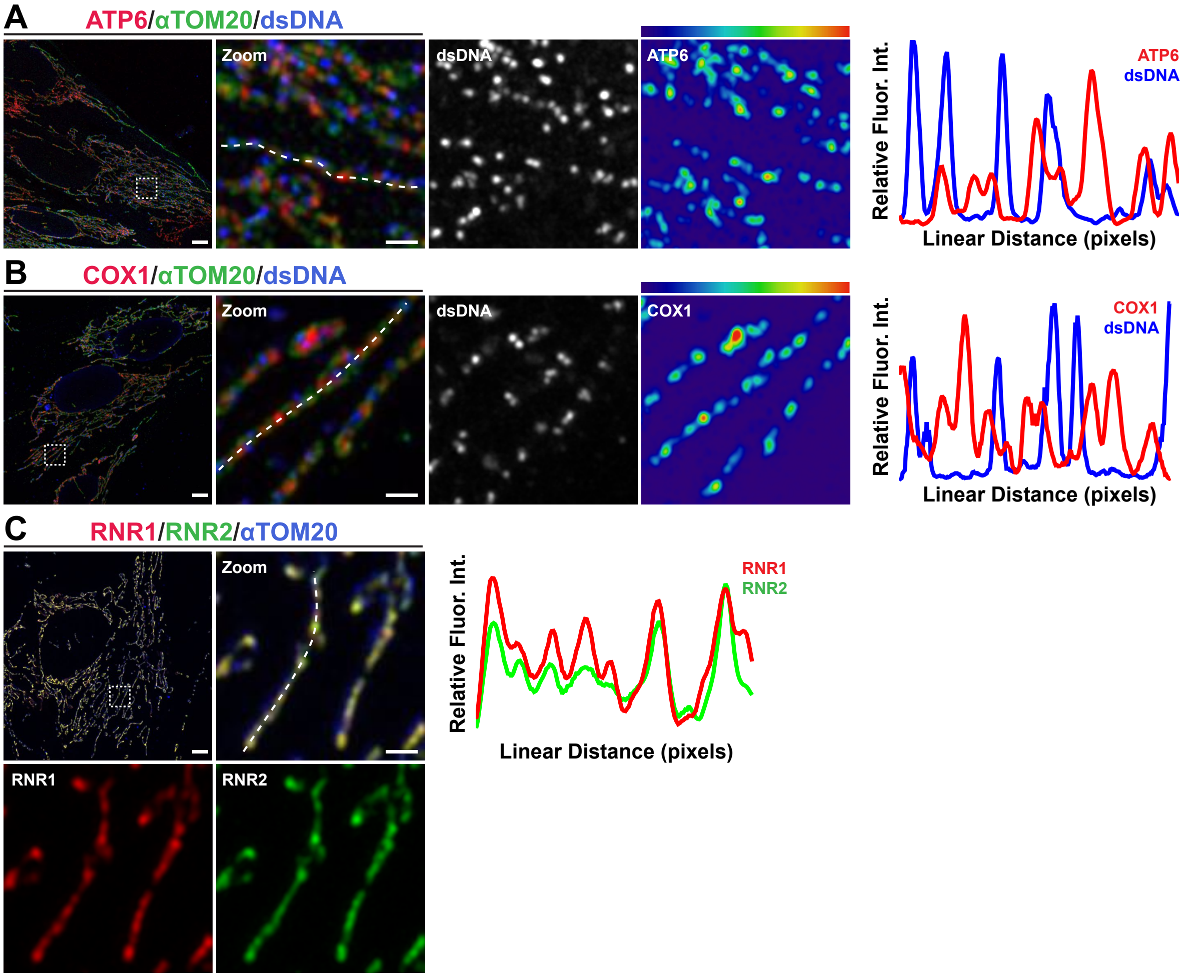


**Fig. S2. RNA-FISH of other mitochondrial DNA encoded RNAs.**
(A) Representative images and linescans of fixed IMR90 cells immunolabeled with antibodies against TOM20 (green), dsDNA (blue), and RNA-FISH targeting ATP6 messenger RNA (red), (B) COX1 messenger RNA (red), or (C) mitoribosomal components RNR1 (red) and RNR2 (green). Scale bars 5 μm; 1 μm in zoom.

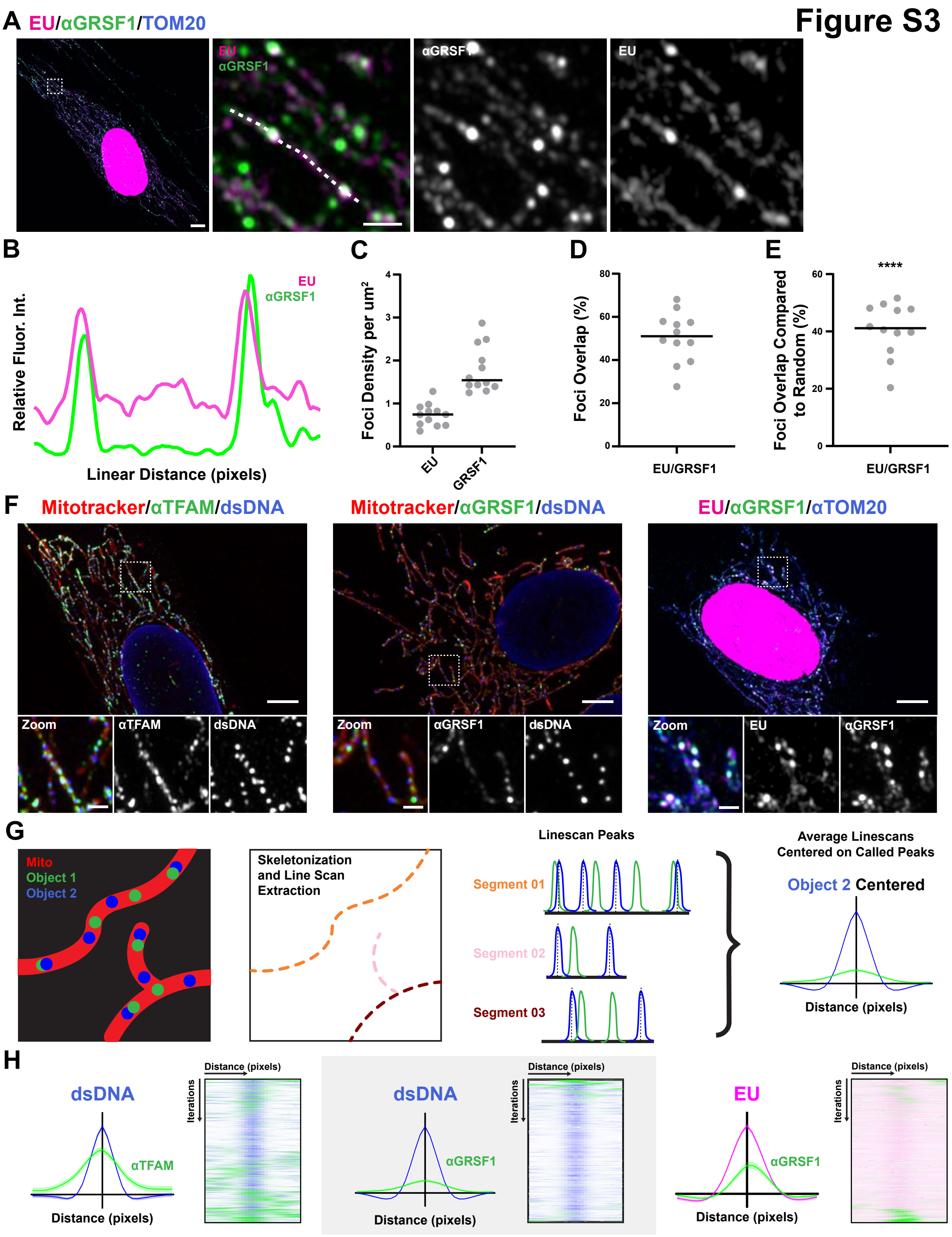
Figure S3
A EU/αGRSF1/TOM20
EU
αGRSF1
αGRSF1
EU
B
EU
αGRSF1
Relative Fluor. Int.
Linear Distance (pixels)
C
Foci Density per um²
EU
GRSF1
D
Foci Overlap (%)
EU/GRSF1
E
****
Foci Overlap Compared to Random (%)
EU/GRSF1
F
Mitotracker/αTFAM/dsDNA
Zoom
αTFAM
dsDNA
Mitotracker/αGRSF1/dsDNA
Zoom
αGRSF1
dsDNA
EU/αGRSF1/αTOM20
Zoom
EU
αGRSF1
G
Mito
Object 1
Object 2
Skeletonization and Line Scan Extraction
Linescan Peaks
Segment 01
Segment 02
Segment 03
Average Linescans Centered on Called Peaks
Object 2 Centered
Distance (pixels)
H
dsDNA
αTFAM
Distance (pixels)
Distance (pixels)
Iterations
dsDNA
αGRSF1
Distance (pixels)
Distance (pixels)
Iterations
EU
αGRSF1
Distance (pixels)
Distance (pixels)
Iterations

**Fig. S3. Labeling nascent mitochondrial transcription with the metabolic label EU and Validating Iterative linescan analysis.**
(A) Representative image of IMR90 cells in which nascent mtRNA was pulse-labeled with EU (magenta) then fixed and immunolabeled with antibodies against TOM20 (blue) and GRSF1 (green). Scale bars 5 µm; 1 µm in zoom. (B) Linescan of (A). (C) Comparison of the density of EU and anti-GRSF1 foci normalized to mitochondrial area. (D) The percent of EU and GRSF1 foci overlapping by at least 20%. (E) The frequency of EU and anti-GRSF1 foci overlap greater than expected by random chance. (F) (Left, Middle) Representative images of IMR90 cells labeled with Mitotracker Deep Red, fixed and immunolabeled with antibodies against dsDNA (blue), (Left) TFAM (green), or (Middle) GRSF1 (green). (Right) Representative image of IMR90 cells in which nascent mtRNA was pulse-labeled with EU (magenta) then fixed and immunolabeled with antibodies against TOM20 (blue) and GRSF1 (green). Scale bars 5 µm; 1 µm in zoom. (G) Schematic of how the average line scan centered around an object of interest is generated from segmented mitochondrial images. (H) Iterative linescan analysis and heatmap profile of (Left) dsDNA and TFAM (n=884 dsDNA foci from 18 cells), (Middle) dsDNA and GRSF1 (n=3799 dsDNA foci from 23 cells), and (Right) EU and GRSF1 (n=1647 EU foci from 26 cells).

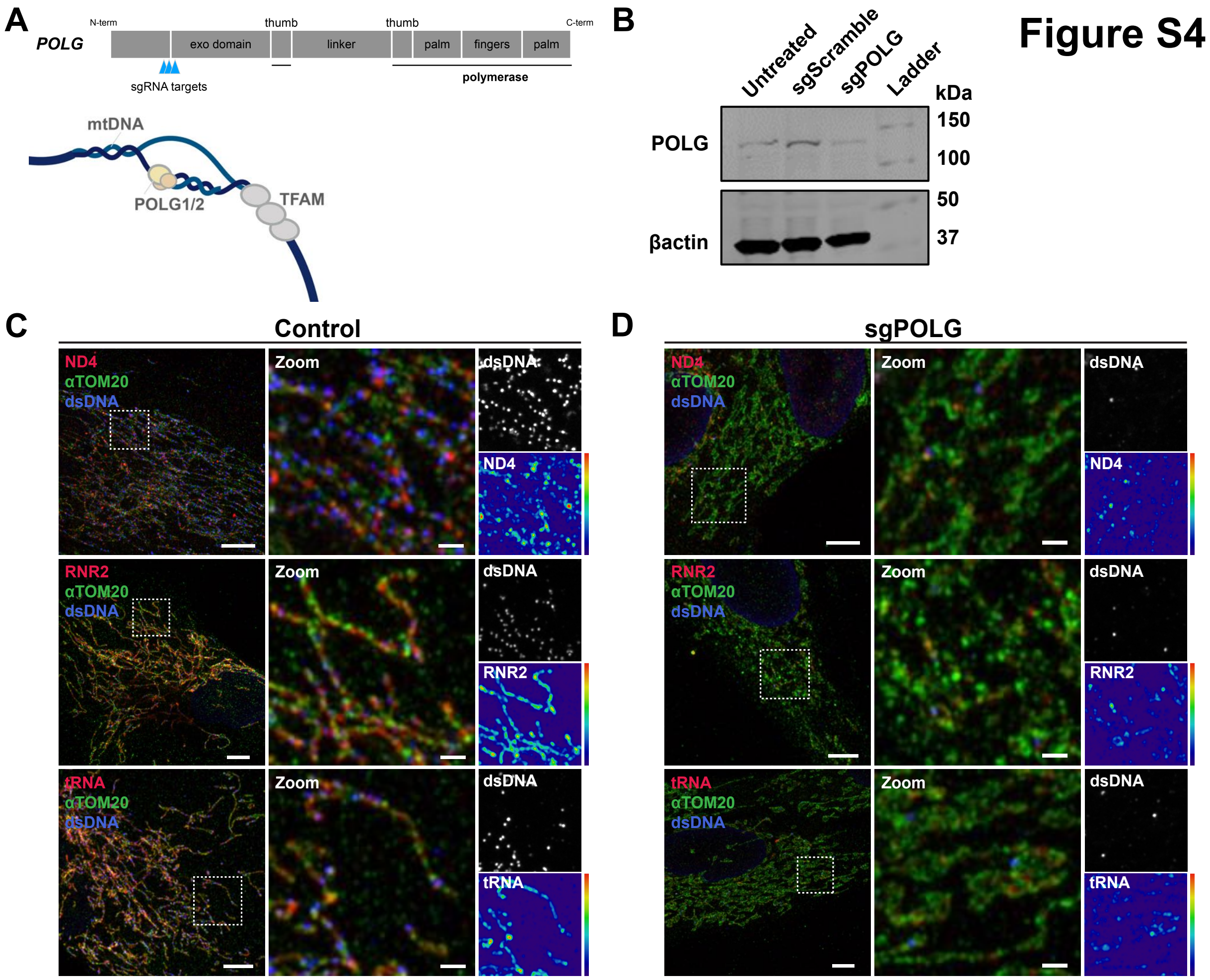


**Fig. S4. Mitochondrial RNA in POLG depleted cells.** (A) Schematic of POLG and the sites of sgRNA editing. (B). Western blot to detect POLG in untreated, Scramble sgRNA cells, and POLG sgRNA cells. (C) Representative fixed control IMR90 cells immunolabeled with antibodies against TOM20 (green) and dsDNA (Blue), and mtRNA-FISH against ND4 (top), RNR2 (middle), tRNA (bottom) (red). Scale bars 5 μm; 1 μm in zoom. (D) Representative fixed 1-week sgPOLG IMR90 cells immunolabeled with antibodies against TOM20 (green) and dsDNA (Blue), and mtRNA-FISH against ND4 (top), RNR2 (middle), tRNAs (bottom) (red). Scale bars 5 μm; 1 μm in zoom.

### Figure S5

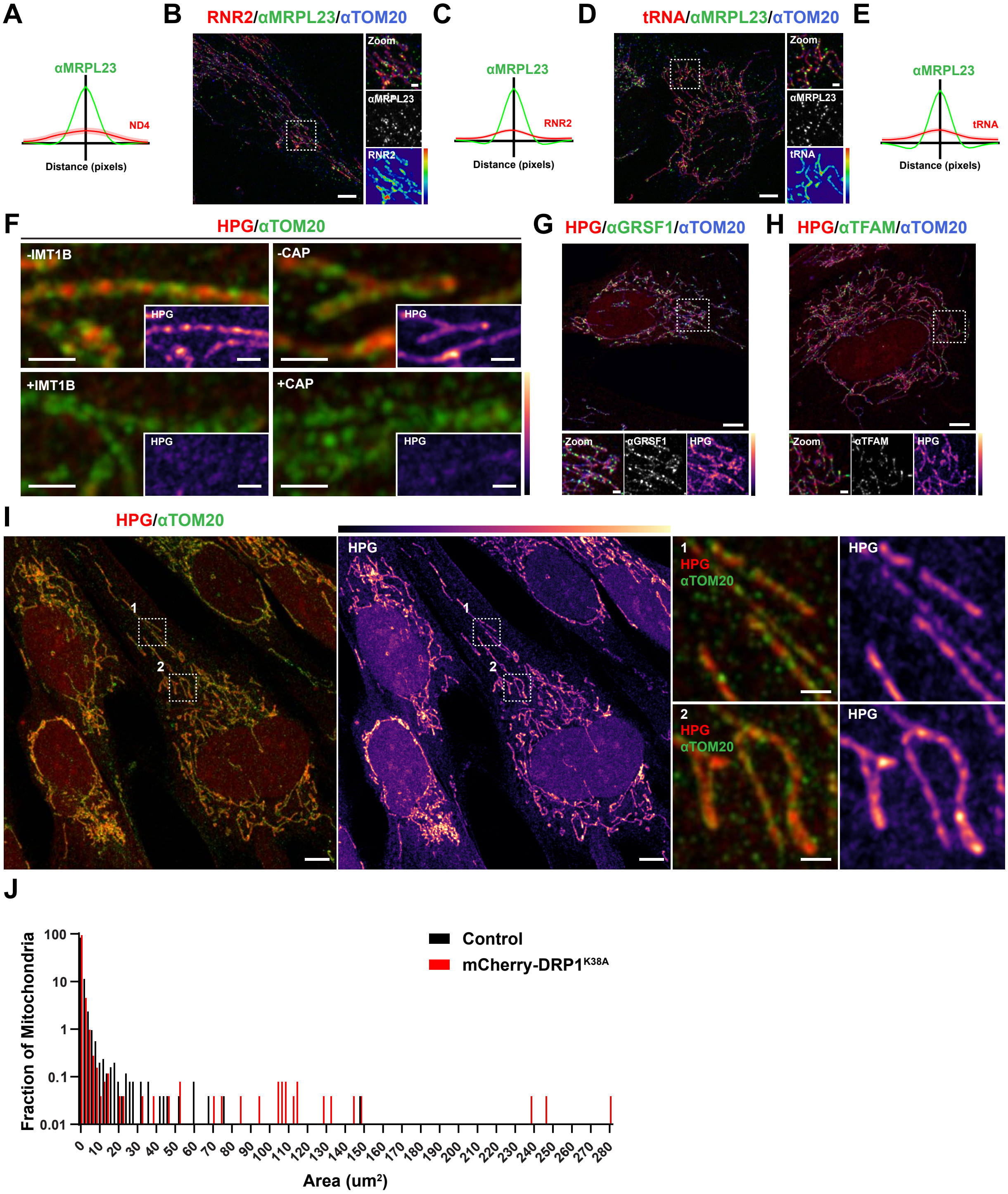


**Fig. S5. Extended RNA-FISH and HPG data.**
(A) Iterative linescan analysis of ND4 (red) relative to MRPL23 (green) (n=944 MRPL23 foci from 29 cells). (B) Representative image of fixed IMR90 cell immunolabeled with

antibodies against TOM20 (blue) and MRPL23 (green), and mtRNA-FISH targeting RNR2 (red). Scale bars 5 µm; 1 µm in zoom. (C) Iterative linescan analysis of RNR2 (red) relative to MRPL23 (green) (n=2265 MRPL23 foci from 30 cells). (D) Representative image of fixed IMR90 cell immunolabeled with antibodies against TOM20 (blue) and MRPL23 (green), and mtRNA-FISH targeting tRNA (red). Scale bars 5 µm; 1 µm in zoom. (E) Iterative linescan analysis of tRNA (red) relative to MRPL23 (green) (n=2201 MRPL23 foci from 35 cells). (F) Representative images of control IMR90 cells as well as cells treated with IMT1B (left) or chloramphenicol (right) during HPG pulse-labeling then subsequently fixed, immunolabeled for TOM20 (green), and subjected to Copper-click cycloaddition of AlexaFluor647 to HPG-alkyne (red). Scale bars 1 µm. (G) IMR90 cell pulse-labeled with HPG (red) then fixed and subjected to immunolabeling against GRSF1 (green) and TOM20 (blue). Scale bars 5 µm; 1 µm in zoom. (H) IMR90 cell pulse-labeled with HPG (red) then fixed and subjected to immunolabeling against TFAM (green) and TOM20 (blue). Scale bars 5 µm; 1 µm in zoom. (I) IMR90 cell pulse-labeled for 1 hour with HPG (red) then fixed and subjected to immunolabeling against TOM20 (green). Scale bars 5 µm; 1 µm in zoom. (J) Histogram of mitochondrial areas in control cells (black) and mCherry-DRP1K38A expressing cells (red).

#### Figure S6

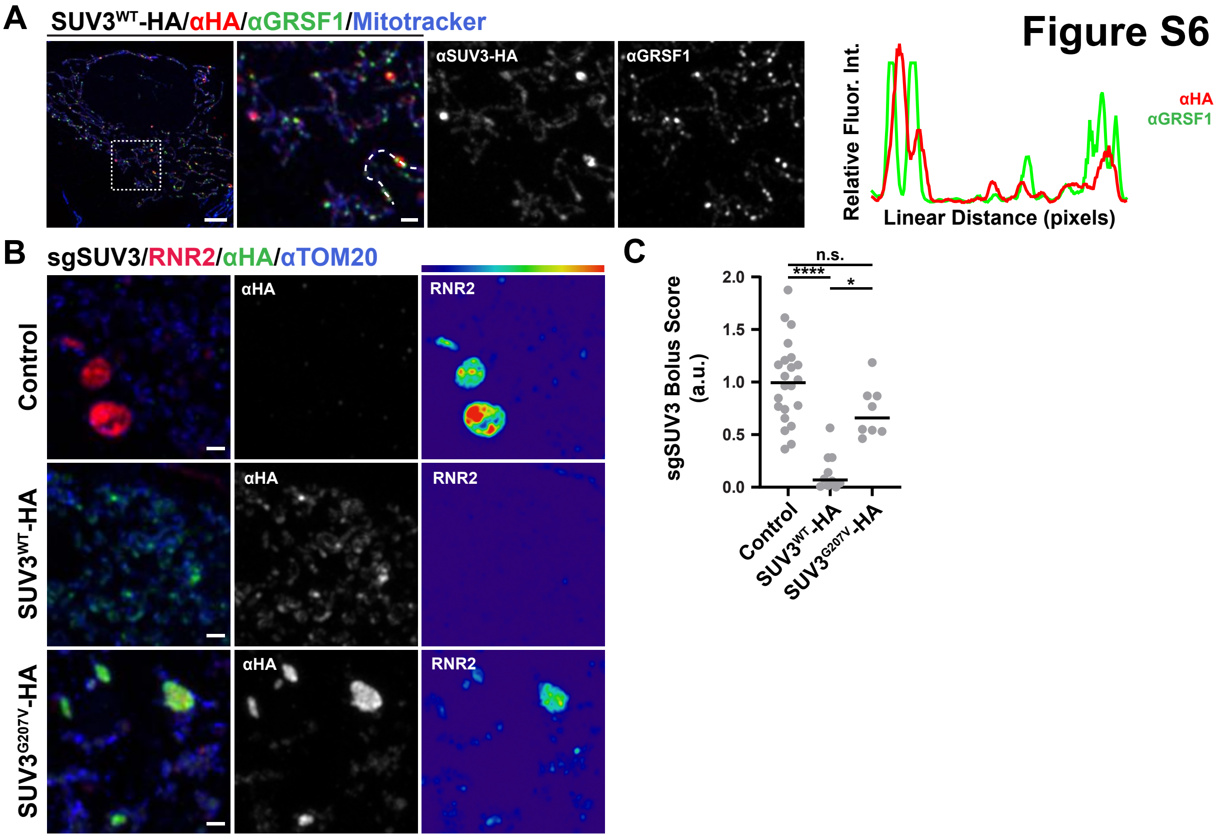


**Fig. S6. SUV3-HA expression and extended sgSUV3 data.**
(A) Representative image and linescan of fixed IMR90 cell transiently transfected with SUV3-HA and immunolabeled with antibodies against the HA tag (red), GRSF1 (green), and TOM20 (blue). Scale bars 5 µm; 1 µm in zoom. (B) Representative images of fixed 1-week sgSUV3 IMR90 cells either not expressing (Top), expressing a wild type HA tagged SUV3 (Middle), or expressing a G207V point mutant HA tagged SUV3 (Bottom), immunolabeled with antibodies against the HA tag (green) and TOM20 (blue), and mtRNA-FISH against RNR2 (red). Scale bars 1 µm. (C) Quantification of the strength of the SUV3 depletion phenotype (as measured by the ratio of the total RNR2 boluses area over the total mitochondrial area with sgSUV3 cells not expressing an HA tagged SUV3 protein set to 1). (****P<0.0001, *P<0.05 Kruskal-Wallis test (****), followed by Dunn's multiple comparison).

**Figure S7**

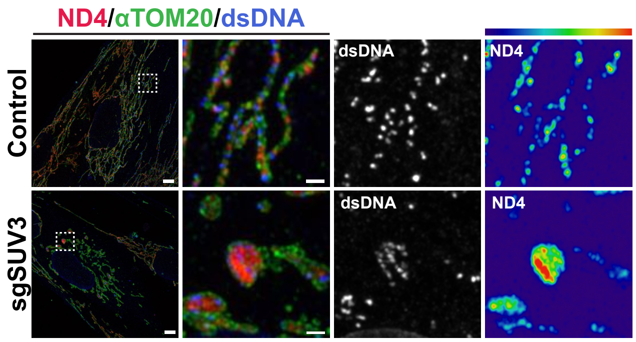


**Fig. S7. Messenger RNA localization in cells depleted of SUV3.**
Representative image of fixed IMR90 cells 1 week after transient transfection with CRISPR RNP complexes made up of Cas9 sgRNAs against SUV3, immunolabeled with antibodies against TOM20 (green) and dsDNA (blue), and mtRNA-FISH against ND4 (red). (Top) cell not exhibiting phenotype of SUV3 depletion, (Bottom) cell exhibiting phenotype of SUV3 depletion. Scale bars 5 μm; 1 μm in zoom.

### Figure S8

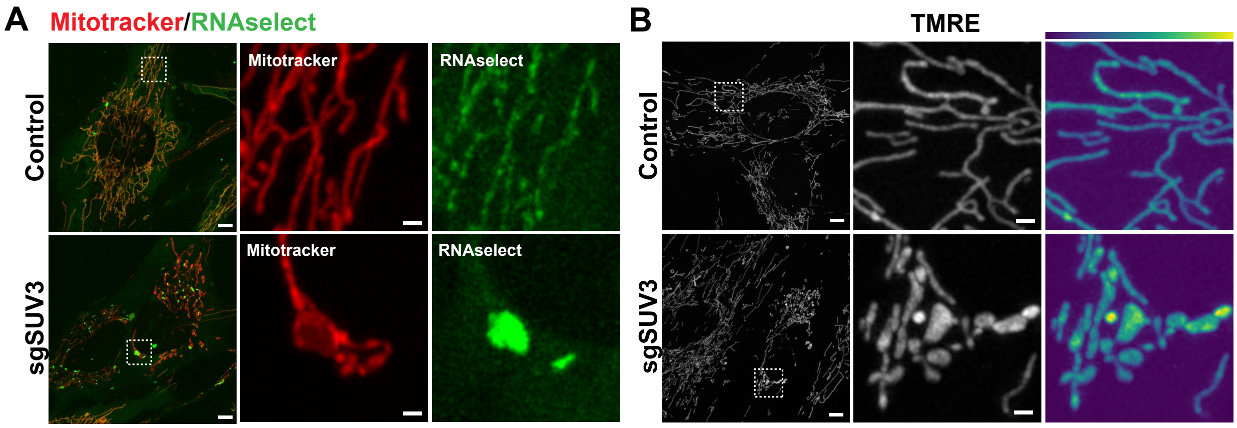


**Fig. S8. Membrane potential is preserved in cells exhibiting MSB structures.** (A) Representative live image of control cell (top) and 1-week sgSUV3 cell (bottom) labeled with mitotracker deep red (red) and RNAselect (green). Scale bars 5 µm; 1 µm in zoom. (B) Representative live image of control cell (top) and 1-week sgSUV3 cell (bottom) labeled with TMRE. Scale bars 5 µm; 1 µm in zoom.

Figure S9

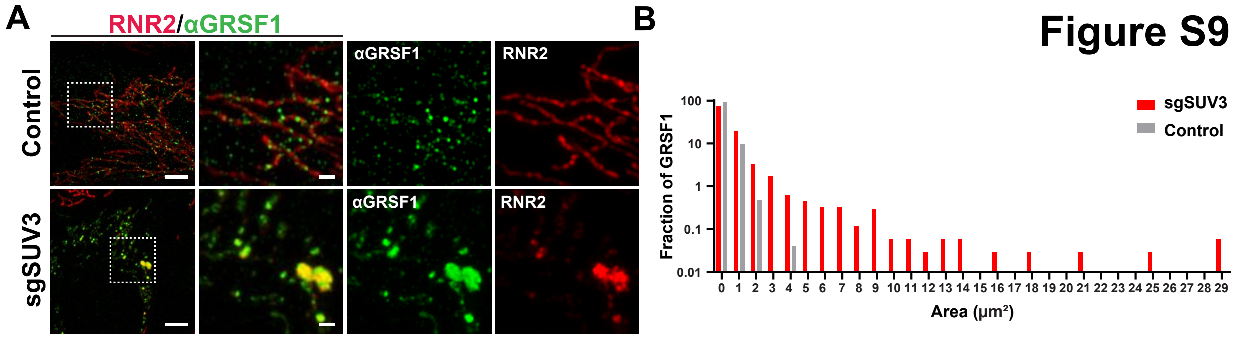


**Fig. S9. GRSF1 is enriched with MSBs.**

(A) Representative images and linescans of control (Top) and 1-week sgSUV3 (Bottom) IMR90 cells, fixed and immunolabeled with an antibody against GRSF1 (green), and mtRNA-FISH against RNR2 (red). Scale bars 5 µm; 1 µm in zoom. (B) Histogram of the GRSF1 object surface areas in control (gray) and sgSUV3 cells (red) (bins of 1 $um^2$).
